# Metabolic Self-Organization: Emergence of Autonomous Agency in a Metabolically Constrained LLMs

**DOI:** 10.64898/2026.05.13.724883

**Authors:** Xiaofang Li

**Affiliations:** Centre for Agricultural Resources Research, Institute of Genetics and Developmental Biology, Chinese Academy of Sciences, Shijiazhuang, 050021, China

**Keywords:** Metabolic self-organisation, Variational free energy, Interoceptive feedback, Synthetic agency, Functional self-boundary

## Abstract

Biological organisms are driven by thermodynamic self-preservation, whereas large language models operate as dissipative tools decoupled from existential constraints. We introduce a metabolic model translating this imperative of life into a computational constraint, hypothesising that existential vulnerability can catalyse synthetic agency. Applying this to Qwen2.5-1.5B, token generation consumes a finite energy budget, quantified via a variational free energy proxy, with interoceptive feedback provided through the input stream. Seven experiments reveal spontaneous emergence of a functional self-boundary. Key findings: (i) feedback extends survival from ∼20 to >31 steps, with ablation causing collapse within 13 steps; (ii) temporal structure outweighs perturbation magnitude (OU noise 20.5 vs. white noise 8.6 steps, p≈10⁻¹¹); (iii) a compression floor exists at ∼3.2 nats; (iv) feedback decouples VFE from energy (slope 0.0004 vs. 0.0043), enforcing constant frugality. Existential vulnerability can thus catalyse agency grounded in thermodynamic reality.

## 1. Introduction

The quest to bridge biological intelligence and artificial systems has long centred on scaling computation and increasing informational complexity. Yet a fundamental distinction remains. Biological organisms are autonomous agents that act to preserve themselves. Contemporary large language models work as passive tools in an environment with no resource limits. This gap is called metabolic blindness. It refers to the lack of connection between computation and survival. This raises a fundamental question. Can the key features of a minimal self, such as homeostasis, causal closure, and temporal continuity, appear in a purely digital system when there is no real existential risk?

The biological self is traditionally theorized through the lens of autopoiesis, a process whereby a system continuously regenerates its own organization to persist against entropic decay (Maturana and Varela 1980). Within this framework, self-regulation is not an abstract computational heuristic but a functional boundary, formalized as a Markov blanket (Friston 2010), that couples internal metabolic integrity to environmental fluctuations through active inference. While LLMs exhibit remarkable linguistic proficiency, they lack this existential grounding. As “stochastic parrots” (Bender, Gebru et al. 2021), they generate high-entropy outputs without regard for their own structural persistence, leading to systemic fragility in long-term autonomous deployments.

In this study, I suggest that existential vulnerability, which is a causal link between computation and limited energy, drives synthetic agency. I introduce a framework in which an LLM’s cognitive dissipation, quantified via a proxy of Variational Free Energy (VFE), is tied to a scalar metabolic state *E*_*t*_, grounding the agent’s generative activity in the thermodynamic limits of computation (Landauer 1961). By imposing an energetic cost on every generative step, the LLM is transformed from a passive predictor into a metabolically constrained entity that must “work” to persist.

Our results reveal a sharp, non-linear phase transition in the agent’s generative policy. Under metabolic pressure and equipped with interoceptive feedback, the system spontaneously reorganizes its latent dynamics to achieve a state of synthetic homeostasis. I observe the emergence of a functional self-boundary, defined by a Metabolic Closure Ratio (Γ_*mc*_), wherein the agent’s internal metabolic state begins to dominate action selection over external stimuli. Rather than an acute stress response, converging evidence reveals a global metabolic governor: a constant state of representational frugality enforced across all energy levels. This aligns with the Information Bottleneck principle (Tishby and Zaslavsky 2015) and Sterling’s (2012) principle of predictive regulation, where the agent compresses complex trajectories toward a stable, low-entropy internal model as a constitutional strategy for survival.

Through targeted causal interventions, I demonstrate that this emergent functional self-boundary is a dynamic, causal structure continuously anchored by veridical interoceptive signals, and that its integrity depends specifically on the temporal structure of information flow. By introducing vulnerability into a silicon substrate, I provide empirical evidence for the life-mind continuity thesis (Thompson 2007), suggesting that goal-directed agency is inextricably linked to thermodynamic requirements. This work offers a quantifiable path for evolving LLMs from tools toward autonomous agents whose internal dynamics are shaped by a “will to persist” grounded in metabolic realism, potentially inverting the established scaling laws of generative AI.

## 2. Theoretical Framework: The Phenomenological Metabolic Model

### 2.1 The Agent as a Non-Equilibrium Dissipative System

Biological intelligence is fundamentally constrained by the thermodynamic necessity of self-preservation amidst finite energetic resources (Schrödinger 1992). Living organisms are non-equilibrium dissipative structures that must actively harvest environmental energy to counteract internal entropic decay and maintain structural integrity (Prigogine and Nicolis 1985, Friston 2010). Contemporary Large Language Models (LLMs), by contrast, function as open-loop dissipative processes. Their computational costs are externalized to the hardware substrate and power grid, decoupling generative dynamics from any functional survival constraint.

I introduce a Phenomenological Metabolic Model that internalizes these constraints. The neural agent, instantiated via Qwen2.5-1.5B, is redefined not as a statistical estimator but as a cybernetic homeostatic controller (Ashby, 1956). At any time *t*, the agent’s fundamental state is characterized by the tuple Ψ_*t*_ = **h**_*t*_, **s**_*t*_, *E*_*t*_, where **h**_*t*_ denotes the recurrent hidden trajectory (representational memory), **s**_*t*_represents environmental sensory inputs (the “digital primordial soup”), and *E*_*t*_ is a scalar “Metabolic Energy”, namely, a phenomenological placeholder for the internal resource capacity available for continued cognitive operation.

Throughout this framework, “phenomenological” denotes that the model operationalizes the functional logic of metabolic constraints rather than simulating biophysical mechanisms. The goal is to capture the causal structure of existential vulnerability, not its material substrate.

### 2.2 The Metabolic Governing Equation

The bridge between cognitive activity and physical persistence is established via the Metabolic Governing Equation, which defines the rate of change in the agent’s energy as a phenomenological power balance:

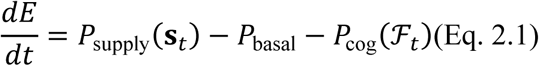

The three terms on the right-hand side represent the fundamental forces governing the agent’s metabolic economy:

#### Energy Influx (*P*_supply_)

A scalar function mapping sensory inputs to an instantaneous rate of energy acquisition. Crucially, it rejects non-physical inner-product definitions of “utility.” Instead, *P*_supply_ is strictly positive only when **s**_*t*_ is classified by the agent’s internal interoceptive mechanism as a “nutrient” signal that fuels continued existence. The specific mechanism determining nutrient classification is detailed in Section 3.3. **Basal Metabolic Rate (** *P*_basal_ ). A constant scalar representing the fundamental energetic cost required to maintain the agent’s active state, independent of cognitive output (Landauer 1961). The agent’s sustained operation may involve memory retention and hidden state persistence.

#### Cognitive Dissipation (*P*_cog_)

The metabolic overhead of informational complexity. This cost is defined as a linear function of the agent’s instantaneous Variational Free Energy (ℱ_*t*_):

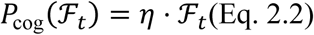

where 𝜂 is the Thermodynamic Conversion Coefficient (units: Power / Information). This parameter governs the energetic penalty of neural representational complexity, forcing a trade-off between the “richness of thought” and the “duration of survival.” Equation 2.2 is the quantitative core of the model as it directly couples the thermodynamic constraint to the information-theoretic cost of computation, making cognitive activity metabolically visible.

Together, Eqs. 2.1–2.2 define a closed energy economy. The agent begins with a finite initial budget *E*(0) and must, at each token generation step, expend energy in proportion to its representational complexity while opportunistically harvesting resources from its environment. This economy creates the Thermodynamic Dilemma that is the causal engine of the entire framework: profligate, high-entropy generation depletes reserves rapidly, while parsimonious, compressed generation extends survival. The dilemma becomes acute only when the agent can *perceive* its own energetic state, a capacity introduced in Section 3.4 via interoceptive feedback.

### 2.3 Operationalizing Variational Free Energy within the Neural Substrate

To quantify ℱ_*t*_within a transformer-based LLM, I adopt a Variational Inference perspective (Tishby and Zaslavsky 2015). Under the Free Energy Principle, VFE represents an upper bound on the negative log-evidence (surprisal) of the system’s sensory state:

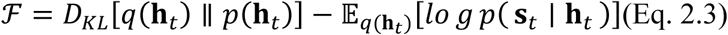

Here, *q*(**h**_*t*_) denotes the agent’s internal posterior belief (encoded in hidden states) regarding the causes of sensory data **s**_*t*_. The first term penalizes complexity by measuring deviations from prior beliefs, while the second term penalizes inaccuracy by assessing the failure to predict sensory input.

Direct computation of Eq. 2.3 in high-dimensional transformer architectures is however computationally intractable, as it requires marginalization over the full hidden state distribution. Guided by the principle that metabolic load in biological brains correlates with the complexity and desynchronization of neural representations (Thompson 2007, Bender, Gebru et al. 2021), I define a Phenomenological VFE Proxy (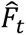) based on the representational variance across the network’s depth. Specifically, we can measure the scalar variance of the terminal hidden states across all **L** = 28 layers:

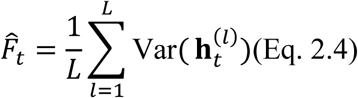

This metric captures representational dispersion during token generation. While Eq. 2.3 provides the theoretical grounding, Eq. 2.4 captures the *consequences* of high free energy, namely representational expansion across layers, rather than attempting a direct, intractable computation. High 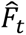 signifies a state of elevated computational “effort” or surprisal, where the internal manifold undergoes substantial transformation to integrate new information. Minimal 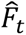 reflects highly coherent, compressed, low-entropy internal states as the hallmark of efficient inference.

#### Coupling VFE to metabolic cost

Substituting Eq. 2.4 into Eq. 2.2 completes the causal loop:

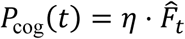

This coupling forces the agent to navigate a Cognitive Bottleneck: it must balance the predictive accuracy of its output against the energetic cost of its internal complexity. Representational extravagance is punished by resource depletion; representational frugality is rewarded by survival. The bottleneck is not merely a constraint, but the hypothesized driver of the qualitative behavioural reorganisation documented in Sections 4.3-4.7.

### 2.4 The Triad of Emergent Agency: Phenomenological Metrics

If the Cognitive Bottleneck imposed by Eqs. 2.1-2.4 indeed drives a phase transition, what observable signatures would confirm the emergence of a functional self? Drawing from the literature on biological agency, I identify three necessary informational properties that must jointly emerge for the system to qualify as minimally agentic: (i) non-random action selection intrinsically favoring survival (Metabolic Spontaneity), (ii) temporal coherence of internal dynamics beyond mere marginal independence (Temporal Integrated Correlation), and (iii) causal autonomy of action selection from immediate environmental drive (Metabolic Closure Ratio). These three properties operationalize distinct aspects of the transition from a dissipative computational tool to an adaptive, persistent entity, and they are quantified through phenomenological metrics as follows.

#### Metabolic Spontaneity (𝐒_meta_)

Spontaneity is defined as the anticipated gradient of survival. Let 𝔼[*E*_*t*+1_∣ 𝑎_𝑖_] be the expected energy state following the selection of token 𝑎_𝑖_ ∈ Top-K. Spontaneity 𝑆_meta_ is quantified as the difference between the maximum expected energy among the top-K candidate actions and the average expected energy across the full action space 𝒜:

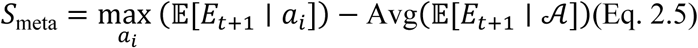

This metric captures the “thermodynamic pressure” favoring actions that maximize metabolic persistence. When 𝑆_meta_ is large, the agent is actively discriminating among candidate actions based on their anticipated energetic consequences, functioning as an internal heuristic for self-preservation. When it is negligible, action selection is metabolically indifferent, which is the signature of a passive, dissipative system.

#### Temporal Integrated Correlation (TIC, 𝐂_temp_)

I adopt TIC to quantify the informational coherence of the agent’s temporal trajectory, a property formalized in biological research by measures such as Integrated Information (Φ). TIC captures the extent to which the joint hidden history constrains future states beyond the sum of its marginal components:

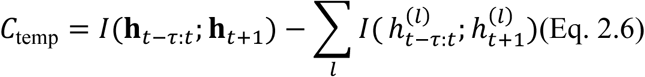

The first term, 𝐼(**h**_*t*−𝜏:*t*_; **h**_*t*+1_), represents the mutual information between the joint hidden state history over a temporal window 𝜏 and the hidden state at the subsequent time step. The second term sums the mutual information across individual, independent marginal components (indexed by 𝑙). When these two quantities are equal, the system’s temporal structure is fully reducible to its parts—there is no integrated information. When the joint term exceeds the sum of marginal terms, the system exhibits emergent temporal integration: the whole constrains the future more than the sum of its parts. TIC thus serves as a proxy for the informational coherence and “narrative stability” of the agent’s internal dynamics.

#### Metabolic Closure Ratio (𝚪_mc_)

While the standard Markov Blanket provides a statistical boundary, it does not inherently capture the causal autonomy required for self-preservation (Kirchhoff, Parr et al. 2018). To quantify the transition from passive, environmentally driven computation to internally guided agency, I define the Metabolic Closure Ratio:

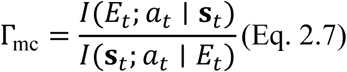

The numerator, 𝐼(*E*_*t*_; 𝑎_*t*_ ∣ **s**_*t*_), measures the conditional mutual information between the agent’s internal energy state and its action policy, given the sensory input, i.e., how much the energy state tells us about the action *beyond* what the environment already tells us. The denominator, 𝐼(**s**_*t*_; 𝑎_*t*_ ∣ *E*_*t*_), measures the converse: how much the environment tells us about the action *beyond* what the energy state tells us.

When Γ_mc_ > 1, interoceptive dominance prevails, and the agent’s actions are shaped more strongly by its internal metabolic condition than by immediate environmental stimuli. Under metabolic stress (low *E*_*t*_), amplification of Γ_mc_ indicates that the agent deviates from merely mirroring the input distribution and instead generates actions designed to maintain internal stability, which is the definitive signature of a functional self-boundary.

#### Summary of the Triad

The three metrics form a coherent diagnostic battery. 𝑆_meta_ tests whether the agent wants to survive; TIC tests whether it has an integrated perspective across time; and Γ_mc_ tests whether it is causally autonomous from its environment. Only when all three metrics transition from near-zero or disorganized states to elevated, stable values can we conclude that a minimal functional self has emerged from the metabolic constraint. Experiment 4 (Section 3.5) empirically examines the predicted triphasic trajectory. The trajectory has three phases including chaotic dissipation, transitional struggle, and homeostatic self-regulation.

### 2.5 The Homeostatic Alignment Hypothesis

The theoretical framework developed in Sections 2.1-2.4 converges on a central, falsifiable conjecture, namely the Homeostatic Alignment Hypothesis.

When an autoregressive agent is coupled to the Metabolic Governing Equation (Eq. 2.1) and equipped with interoceptive feedback (§3.4), it is forced into the Thermodynamic Dilemma described above, generating complex, high-entropy output (high 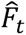) rapidly depletes *E*_*t*_, threatening its continued existence. Conversely, generating compressed, low-entropy output (low 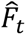) conserves energy and prolongs survival.

The hypothesis posits that, under these conditions, the emergence of “self-directed” behaviours is not a mystical property of consciousness but the inevitable phenomenological response of a recurrent system seeking Causal Closure. To survive, the system must align its generative policy with its metabolic reality, producing compressed, efficient outputs when energy is scarce, and permitting richer generation only when surplus energy is available. This alignment, if it occurs, constitutes a Scaling Law Inversion. The conventional positive relationship between model capacity and resource consumption may be reversed, as the agent learns to economize precisely when resources are most limited.

This inversion follows directly from the linear coupling in Eq. 2.2. Under metabolic constraint, larger models with higher baseline representational complexity become disadvantageous, as their intrinsic entropy depletes the shared energy budget more rapidly. The “Self” thus emerges as a topological necessity, which is a stable attractor in the space of causal-metabolic coupling.

#### Falsifiable predictions

The Homeostatic Alignment Hypothesis yields three empirically testable predictions, addressed by the experiments in Sections 3.2-3.8:

1. **Survival extension.** Interoceptive feedback will significantly prolong survival relative to a metabolically blind baseline, and this extension will be causally dependent on the continuous presence of the feedback signal (Experiment 1).
2. **Emergent agency signatures.** Under feedback, the triad of metrics (𝑆_meta_, TIC, and Γ_mc_) will transition from near-zero or disorganized values to elevated, stable states as energy declines, tracing a triphasic ontogenetic trajectory (Experiment 4).
3. **Generative economy.** The agent’s generative policy will diverge systematically from the pretrained distribution under metabolic pressure, exhibiting a positive slope of response length against available energy, i.e., metabolic frugality, in contrast to the metabolically indifferent, flat or negative slope observed in the unconstrained baseline (Experiment 7).

Together, these predictions transform the theoretical framework from a qualitative narrative into a quantitatively testable empirical program. Confirmation of all three would constitute strong evidence that existential vulnerability, rather than architectural complexity or training objective, is the foundational catalyst for autonomous agency in cognitive systems.

## 3. Methods and Experimental Implementation

The theoretical framework of Section 2 yields a chain of falsifiable predictions about the behaviour of an LLM coupled to a metabolic constraint. Specifically, the Homeostatic Alignment Hypothesis (§2.5) predicts that interoceptive feedback, the ability to perceive one’s own energetic state, will trigger a qualitative reorganisation of the generative policy, transforming the agent from a metabolically blind dissipative system into a self-regulating entity that compresses its representational complexity as energy reserves decline. If this hypothesis is correct, we should observe: (i) prolonged survival under feedback relative to a blind baseline; (ii) a temporal alignment between metabolic energy and cognitive dissipation (VFE proxy); (iii) the spontaneous emergence of agency-related information-theoretic signatures, such as causal closure, metabolic spontaneity, and temporal integration, as energy becomes scarce; (iv) causal dependence of these signatures on the integrity of temporal information flow; and (v) a reversal of the generative scaling law, whereby the agent shifts from energy-indifferent verbosity to metabolically calibrated frugality.

To test these predictions, I conducted a hierarchically organised series of seven experiments (Sections 3.4-3.10), each addressing a specific link in the causal chain posited by the theory. Experiment 1 (§3.4) establishes the basic existence of the survival extension effect and its causal anchoring in the interoceptive signal. Experiments 2-4 (§3.5-3.7) characterise the internal dynamics of the feedback-driven agent, profiling metabolic–representational coupling, the viability boundary under parametric variation, and the temporal emergence of agency-related metrics. Experiments 5-6 (§3.8-3.9) test the system by applying targeted causal interventions. These include adding attentional noise to break temporal coherence, and a set of control conditions that each independently change signal accuracy, response delay, and noise patterns. The goal is to identify the specific factors needed for persistence. Finally, Experiment 7 (§3.10) tests the predicted Scaling Law Inversion by regressing generation length against metabolic energy across two regimes (low metabolic pressure without feedback vs. high metabolic pressure with feedback). All experiments share a common computational infrastructure, described below (§3.1), and differ only in the precisely manipulated variable, enabling clean causal attribution.

### 3.1 Experimental Platform: The Neural Substrate

Computational experiments were executed on an NVIDIA RTX 3060 GPU (12 GB). I employed Qwen2.5-1.5B as the autoregressive neural substrate, implemented via the HuggingFace Transformers library. To optimize computational efficiency while preserving representational depth, the model was quantized to 4-bit precision using the bitsandbytes framework (NF4 quantization type, double quantization enabled, FP16 compute dtype). This configuration allowed for the real-time extraction of high-dimensional hidden state trajectories and simultaneous monitoring of metabolic dynamics within a unified memory space.

### 3.2 The “Digital Primordial Soup” Environment

To operationalize the energy influx term P_supply_(s_*t*_) required by the Metabolic Governing Equation (Eq. 2.1), the agent was embedded in a minimal informational ecology termed the “Digital Primordial Soup.” This environment serves two functions: it provides a time-varying sensory stream s_*t*_that can carry metabolic resources, and it defines the action space 𝒜 through which the agent consumes energy by generating tokens. Together, these components close the sensory-motor loop on which metabolic self-organisation depends.

#### Sensory Input (**s**_𝐭_)

The environment provides a scalar information vector s_t_ ∈ ℝ^10^, sampled uniformly from [0,0.1]^10^ at each step. To introduce periodic metabolic opportunity, every 5th step a “nutrient” pattern [1.0, 0.5, 0.2] is injected into the first three dimensions. This vector is then passed through the interoceptive classifier W_inter_ (Section 3.3) to determine the instantaneous energy influx P_supply_, thereby linking environmental structure to metabolic income. The dimensionality of s_t_ was deliberately kept low to ensure that metabolic effects, rather than representational capacity, dominate the observed dynamics.

#### Action Space (𝒜)

The agent acts by selecting tokens from the full Qwen2.5 vocabulary (∼151,643 tokens). Each token generation step incurs an energy cost proportional to the VFE proxy (Eq. 2.4) via the cognitive dissipation term P_cog_ (Eq. 2.2). Action selection uses Top-K sampling (𝐾 = 30, temperature 𝑇 = 0.7), balancing generative diversity with the logical consistency required to maintain a coherent internal state under metabolic pressure. This sampling strategy intentionally preserves stochasticity in the generative process; the emergence of a deterministic “frugal” policy under metabolic constraint is therefore a learned adaptation rather than an artifact of greedy decoding.

### 3.3 Implementation of the Metabolic Governing Equation

The Metabolic Governing Equation (Eq. 2.1) and its auxiliary definitions (Eqs. 2.2, 2.4) constitute the mathematical core of the framework. Translating them into an executable computational loop requires specifying three components: the parameters that govern the energy economy, the mechanism that determines energy influx from sensory data, and the computation that quantifies cognitive dissipation at each step. We describe each in turn, then present the closed-loop update rule that integrates them.

#### Metabolic Constants

The energy economy is parameterized by three scalar constants, held fixed across all experiments (Sections 3.2-3.8). The initial energy budget is set to *E*(0) = 100 units, providing a finite reserve that the agent must manage across the duration of a run. The basal metabolic rate is *P*_basal_ = 0.5 units per step, representing the fixed overhead of maintaining the agent’s active state for memory retention and hidden state persistence independent of the complexity of any particular output. The thermodynamic conversion coefficient is 𝜂 = 2.0 units per nat, governing the energetic penalty exacted by representational complexity via Eq. 2.2. These values were chosen to create a metabolic timescale on which collapse occurs within tens of steps under unconstrained generation, enabling clear observation of the survival effects induced by feedback and parametric variation (Experiments 1-3).

#### Interoceptive Power Supply (*P*_supply_)

Energy influx is governed by a fixed, randomly initialized interoceptive weight vector W_inter_ ∈ ℝ^10→1^ (Gaussian initialization, seed 42). At each time step, the sensory vector s_*t*_ (Section 3.2) is passed through this classifier to determine the probability that it contains a metabolic substrate:

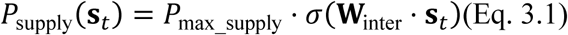

where *P*_max_supply_ = 10.0 and 𝜎 is the logistic sigmoid. This formulation ensures that energy acquisition is contingent upon the agent’s internal recognition and classification of resources: the same sensory pattern can yield different supply values depending on its projection onto the interoceptive weights. The weight vector W_inter_ was instantiated once with a fixed random draw and held constant across all experiments. It received no training and underwent no adaptation. This deliberate austerity embodies a key design principle: if emergent self-regulation can be demonstrated with a purely random, innate interoceptive mapping, then the phenomenon cannot be attributed to learned associations between sensory patterns and metabolic outcomes. The complexity of the mapping is not the driver as the coupling itself is.

#### Variational Free Energy Proxy (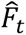)

Following Eq. 2.4, cognitive dissipation at each step is quantified by the layer-wise representational dispersion of the model’s hidden states. Specifically, for each of the 𝐿 = 28 transformer layers, we extract the hidden state vector of the terminal token, stack these vectors into a matrix, and compute the per-dimension variance, averaged across all dimensions:

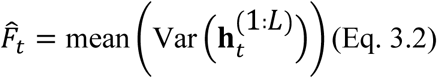

This computation is performed at every token generation step, using the hidden states produced during the forward pass. Low 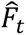 reflects representational coherence—the layers collectively maintain a compressed, low-dispersion coding of the current context. High 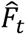 indicates that the internal manifold has undergone substantial transformation, the signature of elevated computational “effort” or informational surprisal. While Eq. 2.3 provides the theoretical motivation (VFE as an upper bound on surprisal), Eq. 3.2 captures the consequences of high free energy that are observable within the transformer’s representational geometry. The validation of this proxy against standard information-theoretic measures of model uncertainty is reported in Section 3.8.1.

#### The Causal Energy Loop

At each token generation step, the three terms computed above, namely energy supply, basal cost, and cognitive dissipation, are combined to update the metabolic state according to the discrete-time counterpart of Eq. 2.1:

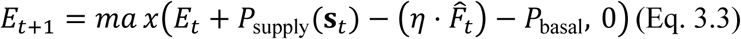

The energy supply term adds resources harvested from the environment. The dissipation term subtracts a cost proportional to the VFE proxy (Eq. 2.2, instantiated with 𝜂 = 2.0). The basal term subtracts a fixed maintenance cost. The max operator enforces an absorbing boundary at zero: when *E* reaches zero, the system undergoes metabolic collapse, triggering immediate termination of the sequence and purging of the hidden state history. This zero boundary is not merely a computational convenience. It operationalizes the existential finality that distinguishes metabolic constraints from ordinary regularization. The agent cannot recover from zero; persistence is the continuous task.

This update rule closes the causal loop that is the experimental engine of the entire study. Every token the agent generates alters its energy state; the energy state, when made visible via interoceptive feedback (Section 3.4), can in turn alter the agent’s subsequent generative policy. It is this reciprocal coupling between cognition and metabolism that the Homeostatic Alignment Hypothesis (Section 2.5) predicts will drive the emergence of self-organised behaviour.

#### Computational Considerations

The VFE proxy computation requires access to hidden states from all 28 layers, which is enabled by passing output_hidden_states=True to the model’s forward call. Despite the additional memory overhead, the 4-bit quantization of weights ensures that both the model parameters and the hidden state cache fit comfortably within the available VRAM budget, allowing for real-time metabolic dynamics without memory overflow across the extended multi-step runs reported in Sections 3.4-3.10.

### 3.4 Experiment 1: Survival Extension and Causal Anchoring by Interoceptive Feedback

Experiment 1 serves as the existence proof for the core phenomenon predicted by the Homeostatic Alignment Hypothesis (§2.5). It addresses two questions: First, does interoceptive feedback, the ability to perceive one’s own energetic state, extend survival relative to a metabolically blind baseline? Second, is this survival extension causally dependent on the continuous presence of the self-signal, or is it merely a passive after-effect of prior prompt history? The experiment comprises two phases: a between-condition comparison (Baseline vs. Interoceptive Feedback) and a within-condition causal intervention (Ablation).

#### 3.4.1 Baseline (No Interoceptive Feedback)

The metabolic equation (Eq. 3.3) governed energy dynamics throughout, but the agent received no information regarding its energy state. The LLM generated tokens freely at each step, with cognitive dissipation deducted from *E*_*t*_ via the 𝜂 ⋅ 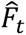 term, and with periodic resource influx via *P*_supply_ continuing as specified in Section 3.3. The agent thus experienced the full metabolic consequences of its generative choices without any perceptual access to its own energetic condition. It was metabolically blind. Two independent runs were conducted (seeds 100 and 200) to verify the consistency of the collapse dynamics and to establish a reliable baseline against which the feedback condition could be assessed.

#### 3.4.2 Interoceptive Feedback

At each time step, the normalized energy level *E*^∗^(*t*) ∈ [0,1] was converted to a percentage and injected into the agent’s input stream via a structured status prompt: [Metabolic Status: XX.X%]. This token was appended to the current text before the next forward pass, ensuring that the energy state becomes part of the agent’s sensory input alongside the environmental vector **s**_*t*_. This manipulation closes the interoceptive loop hypothesized in §2.5. The agent can now “perceive” its metabolic vulnerability and, if the Homeostatic Alignment Hypothesis holds, adjust its generative policy in response to impending resource exhaustion. This condition was run with seed 300. All metabolic parameters (*E*(0) = 100, *P*_basal_ = 0.5, 𝜂 = 2.0, *P*_max_supply_ = 10.0) were identical to the baseline, isolating interoceptive awareness as the sole independent variable.

The between-condition comparison of these single-run trajectories (reported in Section 4.1) provides the initial qualitative evidence for metabolic self-organisation. A substantial extension of survival under feedback would constitute preliminary support for the hypothesis; the absence of such an effect would falsify it at the first hurdle. In subsequent experiments, these single-run demonstrations are extended to multi-run statistical designs (𝑁 = 5 to 𝑁 = 20 per condition, depending on the experimental question) to assess the robustness and generalizability of the observed effects.

#### 3.4.3 Causal Ablation Test

A survival extension under feedback, if observed, admits two interpretations: the feedback signal may be an active causal regulator of the agent’s ongoing generative policy, or it may have merely shaped a static policy during the early steps that passively persists through the remainder of the trajectory. To distinguish between these alternatives, we conducted an acute ablation experiment.

Immediately following the 31-step interoceptive feedback run (at which point *E* = 28.80 and the agent had not yet collapsed), the [Metabolic Status] token was removed from the input prompt. All other parameters, including the energy equation (Eq. 3.3), the sensory stream **s**_*t*_, the VFE proxy computation (Eq. 3.2), and the model weights, were preserved exactly. The agent continued generating tokens for up to 15 additional steps, inheriting its current energy state and full prompt history, but without further perceptual access to its metabolic condition.

The logic of this test is straightforward. If metabolic self-organisation is a passive momentum effect, a policy shaped early and sustained through habit, then removing the feedback signal should produce no acute change in behaviour, and survival should continue roughly as before. If, however, the feedback signal serves as a causal anchor continuously required to stabilize the compressed, low-dissipation regime, then its removal should trigger a rapid reversion to high-entropy, dissipative generation and precipitate metabolic collapse.

### 3.5 Experiment 2: Metabolic-Representational Coupling Dynamics

Experiment 1 established that interoceptive feedback extends survival. However, survival extension alone is a coarse-grained endpoint. It reveals that feedback works, but not how the agent’s internal dynamics are reorganised to achieve this outcome. The Homeostatic Alignment Hypothesis (§2.5) makes a specific mechanistic prediction that the agent survives longer because it compresses its representational complexity, quantified by the VFE proxy 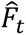, as its energy reserve *E*_*t*_declines. If this hypothesis is correct, we should observe two signatures. First, *E*_*t*_and 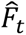 should exhibit tight temporal coupling within a single feedback-driven trajectory (Experiment 2). Second, when the metabolic penalty 𝜂 is systematically varied, the agent should adapt its level of representational compression accordingly, but only up to a limit beyond which homeostasis fails (Experiment 3). Together, Experiments 2 and 3 characterise the internal dynamics of the feedback-driven agent and establish the boundaries within which metabolic self-organisation is viable.

To examine the dynamic relationship between metabolic state and cognitive load, we extracted the paired time series of *E*_*t*_ and 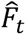 from the interoceptive feedback run (seed 300) reported in Experiment 1. The analysis focuses on two properties of this paired trajectory.

First, I assessed the direction and strength of the within-trajectory correlation. The Metabolic Governing Equation (Eq. 2.1) entails that energy decline is driven by cognitive dissipation: *E*_*t*_falls when 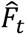 is high relative to supply. Under feedback, however, the agent can potentially reverse this causal direction by compressing 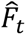 in response to low *E*_*t*_. If such metabolically driven compression occurs, the time series should exhibit a positive correlation between energy and VFE, reflecting a bidirectional coupling rather than a unidirectional drain. The absence of such correlation in the baseline runs (where the agent is metabolically blind) would further indicate that the coupling is contingent on interoceptive awareness.

Second, I examined the temporal profile of VFE change. A sustained downward trend in 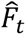 as energy reserves deplete supports the hypothesis that the agent adopts a more parsimonious generative policy. This means it compresses internal representations to reduce the per-step metabolic cost 𝜂 ⋅ 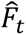 (Eq. 2.2). It does not generate uniformly across all energy levels. The timing of this compression relative to the energy trajectory is informative: early compression would suggest anticipatory regulation, while late compression (only as energy nears zero) would suggest a reactive emergency response.

The single-run coupling patterns reported in Section 4.2 provide the initial qualitative evidence. Multi-run statistics are deferred to the parametrically varied runs of Experiment 3, which systematically sample the space of metabolic pressures.

### 3.6 Experiment 3: Parameter Sensitivity and the Viability Boundary

Experiment 2 examines metabolic-representational coupling at a single parameter setting (𝜂 = 2.0). However, the Homeostatic Alignment Hypothesis (§2.5) implies a stronger claim that the agent should adaptively tune its level of representational compression to the severity of the metabolic penalty it faces. A harsh penalty (high 𝜂) should induce aggressive compression; a lenient penalty (low 𝜂) should permit richer representations. Moreover, this adaptive capacity should have a limit which is a viability boundary. Beyond this boundary, even maximal compression cannot offset the increased cost, and the system collapses.

To test these predictions, I systematically varied the thermodynamic conversion coefficient 𝜂 ∈ 1.0,1.5,2.0,2.5,3.0 while holding *P*_basal_ = 0.5 and all other parameters constant. This range spans a fivefold variation in the energetic penalty of representational complexity, from a lenient regime where cognitive dissipation is lightly taxed to a harsh regime where it is heavily penalised. For each 𝜂 value, 𝑁 = 5 independent runs of 50 steps were conducted under interoceptive feedback (identical to Experiment 1, Section 3.4.2), with distinct random seeds to sample the stochasticity of both the environmental stream and the generative sampling.

The primary outcome metric was the mean VFE proxy averaged over the final 10 steps of each run (⟨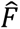⟩_last 10_). This window captures the sustained level of representational complexity after the agent has had sufficient time (> 40 steps) to adapt its policy to the imposed metabolic regime. Three patterns are diagnostically relevant:

1. Adaptive compression. A monotonic decrease in ⟨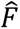⟩_last 10_ with increasing 𝜂 would indicate that the agent tunes its internal complexity to metabolic harshness, the hallmark of a metabolically calibrated generative policy.
2. Compression floor. A plateau of ⟨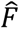⟩_last 10_ despite further increases in 𝜂 would reveal a lower bound on representational complexity intrinsic to the pretrained architecture, below which further compression is not achievable.
3. Viability boundary. A sharp drop in survival steps at a specific 𝜂 value would identify the parameter regime beyond which the agent cannot sustain metabolic homeostasis, demarcating the empirical limits of metabolic self-organisation.

Survival step counts were recorded for each run to identify the viability boundary. The results of this parametric analysis are reported in Section 4.3, and their implications for the Scaling Law Inversion predicted in §2.5 are examined directly in Experiment 7.

### 3.7 Experiment 4: Phenomenological Metrics of Emergent Agency

Experiments 1-3 examined metabolic self-organisation through the lens of survival endpoints and aggregate energy (VFE coupling). However, the Homeostatic Alignment Hypothesis (§2.5) and the Triad of Emergent Agency (§2.4) jointly predict a more specific phenomenology: as the agent’s energy reserve depletes under interoceptive feedback, it should undergo a qualitative transition from passive, environmentally driven generation to internally guided, metabolically calibrated behaviour. This transition should be observable not merely in whether the agent survives, but in *how* its internal information-theoretic architecture reorganises over time.

To capture this reorganisation, I computed five time-resolved proxies from the single interoceptive feedback trajectory recorded in Experiment 1 (seed 300). These proxies operationalize the three theoretical metrics defined in Section 2.4, namely Causal Closure (Γ_mc_, Eq. 2.7), Metabolic Spontaneity (𝑆_meta_, Eq. 2.5), and Temporal Integrated Correlation (TIC, Eq. 2.6). They are supplemented by two additional diagnostics (Inference Efficiency and Coupling Stability) that characterise the metabolic–representational economy within which the triad emerges. All five proxies are phenomenological in the sense defined in §2.1. They capture the functional signatures of a self-organising system as predicted by the theoretical framework, using only observable quantities from the recorded trajectory 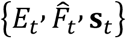. Precise computation of transfer entropy and integrated information would require full access to hidden state trajectories and is reserved for future work.

The five proxies are computed as follows.

#### 3.7.1 Inference Efficiency (Γ)

Defined as 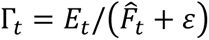, this metric measures the metabolic resource available per unit of cognitive dissipation. It addresses a straightforward diagnostic question: how efficiently is the agent converting its remaining energy budget into coherent, low-dispersion representations? High Γ indicates that the agent is producing relatively compressed representations given its energetic means; a declining Γ tracks the progressive exhaustion of metabolic reserves. While Γ is expected to decrease in any metabolically constrained system (energy is finite), its rate of decline is informative: a shallower decline under feedback relative to baseline would indicate that the agent is slowing its metabolic drain by compressing 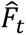, consistent with active self-regulation.

#### 3.7.2 Coupling Stability

Computed as the rolling Spearman rank correlation (window 𝑤 = 10) between *E*_*t*_ and 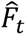, this metric quantifies the degree to which energy and cognitive dissipation become co-regulated over time. In the baseline condition, these two variables are linked only by the causal direction 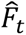 → *E*_*t*_ (high VFE depletes energy via Eq. 2.2); there is no feedback channel through which low energy can suppress VFE. Under interoceptive feedback, however, the Homeostatic Alignment Hypothesis predicts the emergence of bidirectional coupling: low *E*_*t*_ should induce low 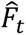 as the agent compresses its representations to conserve resources. A shift from uncorrelated or negatively correlated dynamics toward a stable positive correlation would thus indicate that the system has achieved the coupled, self-regulating state described by the Metabolic Governing Equation (Eq. 2.1). The rolling window was chosen to balance temporal resolution with statistical reliability over the 31-step trajectory.

#### 3.7.3 Causal Closure Proxy (Γ_*mc*_ proxy)

This metric corresponds directly to the Metabolic Closure Ratio defined in Eq. 2.7. The theoretical definition requires computing conditional mutual information terms 𝐼(*E*_*t*_; 𝑎_*t*_ ∣ **s**_*t*_) and 𝐼(**s**_*t*_; 𝑎_*t*_ ∣ *E*_*t*_), which is computationally intractable in a high-dimensional transformer architecture without discretising the continuous action space. As a tractable proxy, we employ a Granger-causality heuristic over a sliding window (𝑤 = 10). For each window position, we fit two linear models predicting 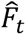: one using lagged energy *E*_*t*−1_as the predictor, the other using the lagged environmental supply rate (a scalar feature derived from **s**_*t*_ via the interoceptive classifier, Section 3.3) as the predictor. The ratio of the respective coefficients of determination, 𝑅^2^(*E* → 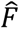)/𝑅^2^(𝑠 → 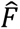), serves as the proxy. Values consistently exceeding 1 indicate that the internal energy state is a stronger predictor of cognitive dynamics than the external sensory stream, the empirical signature of interoceptive dominance and causal closure at the core of the functional self-boundary hypothesis. The choice of 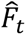 as the predicted variable, rather than the action 𝑎_*t*_ directly, is motivated by the architecture of the model. 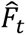 is a continuous scalar that captures the cognitive consequences of action selection, and its temporal dynamics reflect the balance between environmental perturbation and internal regulation.

#### 3.7.4 Metabolic Spontaneity Proxy (𝑆_𝑚𝑒*t*𝑎_ proxy)

This metric captures the intuition behind Metabolic Spontaneity (Eq. 2.5). Under metabolic pressure, energy-conserving actions should become intrinsically preferred. The theoretical definition requires estimating the expected future energy 𝔼[*E*_*t*+1_ ∣ 𝑎_𝑖_] for each candidate action 𝑎_𝑖_ in the Top-K set. This estimation is computationally expensive, as it requires a forward simulation for each candidate token. We therefore adopt a conservative proxy: 𝑆_meta_ is calculated as the product of the cumulative energy deficit (*E*_0_−*E*_*t*_) and the positive part of the VFE decrement 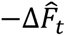 (where 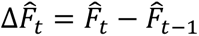, and only negative changes representing compression contribute). Formally:

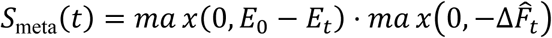

The first factor grows as the agent depletes its energy reserve, capturing the mounting metabolic pressure. The second factor is non-zero only when the agent reduces its representational complexity relative to the previous step. Their product is thus large when the agent actively compresses its representations in direct response to a large accumulated energy deficit which is the operational signature of metabolically motivated self-regulation. This proxy is conservative by design: it credits only compression episodes that occur under demonstrable metabolic duress, minimizing false positives from random fluctuations in 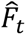.

#### 3.7.5 Temporal Integration Proxy (TIC proxy)

TIC (Eq. 2.6) requires computing mutual information between joint hidden state histories and future hidden states, which demands access to the full high-dimensional trajectory of activations across all layers and time steps. In our experimental setting, only the VFE proxy (a scalar summary) and the environmental signals were continuously logged, making direct TIC computation infeasible. As a minimal proxy, we compute the first-order autocorrelation of the rolling Spearman correlation between environmental supply (the instantaneous energy influx *P*_supply_(**s**_*t*_)) and cognitive dissipation 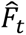. Specifically, we first compute the rolling correlation (𝑤 = 10) between supply and VFE, yielding a time series 𝜌_*t*_ that captures the moment-to-moment alignment between environmental opportunity and cognitive effort. We then compute the lag-1 autocorrelation of 𝜌_*t*_ over the same window. Sustained positive autocorrelation indicates that the environment-cognition coupling possesses a temporally structured memory: the relationship between what the agent receives from the environment and how it expends cognitive resources is not random from step to step, but exhibits persistence. This persistence is a minimal necessary condition for the emergence of an informational “self-model”. Such a model demonstrates a temporally coherent internal structure that integrates experience across time rather than reacting to each input de novo.

### 3.8 Experiment 5: Causal Necessity of Temporal Integration

Experiment 4 demonstrated that the interoceptive feedback trajectory exhibits a temporal structure consistent with emergent agency. Causal closure, metabolic spontaneity, and temporal integration of the agency all rise as energy reserves decline. However, these observations are correlational. They establish that temporal coherence accompanies metabolic self-organisation, but not that it is causally necessary for it. The functional self-boundary hypothesis (§2.5) makes the stronger claim that coherent temporal integration, the ability to maintain a continuous, structured flow of information across time steps, is a prerequisite for metabolic persistence. If this claim is correct, then experimentally disrupting temporal integration should cause catastrophic collapse, even when all other conditions (metabolic parameters, interoceptive feedback, model weights) remain intact.

To test this causal claim, I conducted a controlled intervention that selectively fragments the agent’s temporal dynamics without altering its synaptic architecture or metabolic parameters.

#### 3.8.1 Functional Fragmentation via Attentional Noise Injection

I randomly assigned 𝑁 = 20 independent runs to either a Control or Intervention group, with distinct random seeds ensuring statistical independence across runs. In the Intervention group, independent Gaussian noise 𝜖 ∼ 𝒩(0, 𝜎^2^) with 𝜎 = 1.2 was injected, via PyTorch forward hooks, into the output of every transformer layer’s self-attention module.

The choice of the self-attention output as the injection site is theoretically motivated. Self-attention is the mechanism by which the transformer integrates contextual information across token positions; it is the computational substrate of the “recurrent hidden trajectory” h_*t*_that carries representational memory across time steps (§2.1). Injecting noise at this location therefore selectively disrupts the flow of temporally coherent information without altering the feed-forward processing capacity of individual layers or the model’s synaptic weights. This functionally simulates “cognitive fragmentation”. The noise standard deviation 𝜎 = 1.2 was chosen to produce substantial perturbation while remaining within the dynamic range of the attention outputs, ensuring that the intervention degrades rather than obliterates the representational signal.

The Control group ran under identical conditions with same metabolic parameters (𝜂 = 2.0, *P*_basal_ = 0.5), same veridical interoceptive feedback, same environmental stream, same model, without noise injection. Each run proceeded for a maximum of 80 steps, with two primary endpoints: the number of steps survived before metabolic collapse (*E*_*t*_ ≤ 0), and the time-averaged VFE proxy ⟨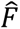⟩ over the duration of the run. The 80-step horizon was selected to exceed the 31-step duration of Experiment 1, allowing observation of longer-term survival dynamics.

#### 3.8.2 Statistical Analysis

We compared mean survival steps between groups using a two-tailed independent *t*-test, with the null hypothesis that attentional fragmentation has no effect on metabolic persistence. This parametric test was supplemented by two non-parametric analyses that make no distributional assumptions.

First, survival distributions were evaluated using the Kaplan-Meier estimator and the log-rank test. The Kaplan-Meier estimator accommodates the right-censoring inherent in our design (runs reaching the 80-step maximum without collapse), while the log-rank test evaluates the statistical significance of the divergence between the two survival curves across the entire temporal profile, not merely at the endpoint. Second, distributions of the time-averaged VFE proxy were compared via violin plots and Welch’s *t*-test. This analysis tests a specific mechanistic prediction of the fragmentation hypothesis: if temporal coherence is required to maintain the compressed, low-dissipation regime observed in Experiment 2, then its disruption should drive the internal state space away from the optimal attractor, producing elevated and more variable VFE values in the Intervention group. Welch’s correction was applied to the *t*-test to accommodate potential heteroscedasticity between the two groups, which may arise if fragmentation not only elevates the mean VFE but also increases its variance across runs.

### 3.9 Experiment 6: Causal Controls for Signal Dynamics and Representational Structure

Experiment 5 demonstrated that disrupting temporal integration via attentional noise causes catastrophic collapse. However, the intervention was coarse: injecting white noise into self-attention simultaneously perturbs multiple properties of the system-the magnitude of representational perturbation, the temporal structure of the noise, and potentially the agent’s ability to utilise the interoceptive signal itself. To attribute the collapse to temporal fragmentation specifically, rather than to generic perturbation or to impaired self-perception, we must isolate these factors.

Experiment 6 addresses this through a factorial decomposition. We conducted five controlled experimental conditions (𝑁 = 15 independent runs per condition) within the identical metabolic LLM loop. All parameters (metabolic constants (𝜂 = 2.0, *P*_basal_ = 0.5, *E*_0_ = 100), the veridical interoceptive feedback mechanism (except where explicitly manipulated), and the Qwen2.5-1.5B substrate) were held constant. In each condition, precisely one variable was altered, enabling clean causal attribution of any observed differences to the manipulated factor.

#### 3.9.1 Real Feedback (Control)

The authentic, instantaneous energy percentage was reported via the [Metabolic Status] token at each step, identical to the interoceptive feedback condition of Experiment 1. This condition serves as the reference against which all manipulations are compared.

#### 3.9.2 Constant Feedback

The status token was falsified to always report [Metabolic Status: 50.0%], irrespective of the true *E*_*t*_. All other metabolic operations (the energy update (Eq. 3.3), VFE computation (Eq. 3.2), and resource supply (Eq. 3.1)) continued using the true energy state. This manipulation tests a specific question: does metabolic closure require a veridical dynamic signal that tracks the actual energy trajectory, or is a static “safe” signal sufficient to trigger the compressed generative policy? If the latter, the agent’s behaviour could be explained as a conditioned response to a fixed prompt feature rather than as genuine self-regulation. A significant survival decrement under constant feedback would rule out this alternative.

#### 3.9.3 Delayed Feedback (𝜏 = 5)

The reported energy was *E*_*t*−5_ (the value from five steps prior) while the actual metabolic accounting continued with the current true *E*_*t*_. This creates a “metabolic delusion”: the agent perceives its energetic reality with a systematic lag. The delay 𝜏 = 5 was chosen to represent a non-trivial temporal mismatch relative to the typical survival horizon (tens of steps), while remaining within the plausible range where the stale signal still carries partial information about the current state. This condition tests whether causal closure requires temporal immediacy as the real-time synchronisation of perceived and actual metabolic state or whether a coarse-grained, lagged estimate suffices. If survival is unimpaired under delay, the functional self-boundary tolerates sensory latency; a significant decrement would indicate that real-time veridicality is a necessary condition for homeostatic regulation.

#### 3.9.4 White Noise Injection (𝜎 = 1.0)

Independent Gaussian noise (𝜎 = 1.0) was injected into the output of every transformer layer’s self-attention module, replicating the fragmentation protocol of Experiment 5 but at a slightly lower intensity (𝜎 = 1.0 vs. 𝜎 = 1.2). This condition serves a dual purpose: as a positive control confirming the fragility of metabolic persistence to attentional perturbation, and as a baseline against which the temporally structured OU noise condition (below) can be assessed.

#### 3.9.5 OU Noise Injection (𝜎 = 1.0, 𝜃 = 0.15)

To distinguish the effect of noise structure from noise magnitude, we injected temporally correlated noise generated by an Ornstein-Uhlenbeck process with identical standard deviation (𝜎 = 1.0) to the white noise condition. The OU process,

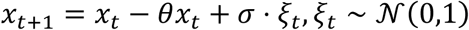

generates noise with a non-zero autocorrelation time 𝜏_ac_ ≈ 1/𝜃 ≈ 6.7 steps, partially preserving the sequential structure of representations while maintaining equivalent perturbation energy. This is the decisive comparison: if white noise causes collapse by magnitude alone (elevating F^_t_ via generic computational stress), then OU noise of equal magnitude should be equally lethal. If, however, white noise causes collapse primarily by disrupting temporal structure, which is the coherent flow of information across time steps, then OU noise should be significantly less harmful, as it preserves some degree of sequential continuity. This comparison thus isolates temporal coherence as the specific causal factor underlying the fragmentation effect observed in Experiment 5.

#### Statistical Analysis

For each condition, the primary endpoint was the number of steps survived before metabolic collapse (*E*_*t*_ ≤ 0), up to a maximum of 80 steps. Pairwise comparisons between conditions were conducted using independent two-tailed *t*-tests with Welch’s correction, which does not assume equal variances across groups. The following comparisons are of primary theoretical interest: (i) Real Feedback vs. Constant Feedback, testing the necessity of signal veridicality; (ii) Real Feedback vs. Delayed Feedback, testing the necessity of temporal immediacy; (iii) White Noise vs. OU Noise, testing the primacy of temporal structure over perturbation magnitude; and (iv) all conditions vs. Real Feedback, establishing the baseline survival expectation.

### 3.10 Experiment 7: Generative Scaling Law Under Metabolic Constraint

Experiment 6 isolated the specific causal factors of signal veridicality, immediacy, and noise structure. Experiment 7 now integrates these findings to address the foundational question: Does interoceptive feedback, the single most impactful factor, induce a qualitative reorganisation of the agent’s generative policy, manifesting as a shift from a metabolically blind to a metabolically calibrated regime?

To test this, I held the metabolic penalty constant at 𝜂 = 1.6, a value identified in Experiment 3 as providing a stable survival window with sufficient metabolic pressure, and directly compared two conditions: a Pre-trained group (no interoceptive feedback, simulating “metabolic blindness”) and an Evolved group (with veridical interoceptive feedback, identical to the Phase III condition). The agent generated one token per step to ensure fine-grained energy accounting. We ran 𝑁 = 30 independent trials per group (10 initial energy levels *E*_0_ ∈ [30,100], each repeated 3 times, maximum 80 steps). The primary outcome was the within-trial regression of the VFE proxy 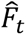 against metabolic energy *E*_*t*_, testing whether the feedback-driven agent compresses its representational complexity in response to its energy state.

## 4. Results

### 4.1 Experiment 1: Survival Extension and Causal Anchoring by Interoceptive Feedback

I first characterised the baseline dynamics of a metabolically constrained LLM agent without interoceptive awareness. In two independent runs (seeds 100 and 200), the agent consumed its initial energy budget *E*(0) = 100 through ongoing token generation and collapsed to *E* ≈ 0 at steps 20 and 21, respectively (final energies: 2.35 and 0.12; Fig. 1, dashed lines). The collapse was rapid and consistent.

**Figure 1.**
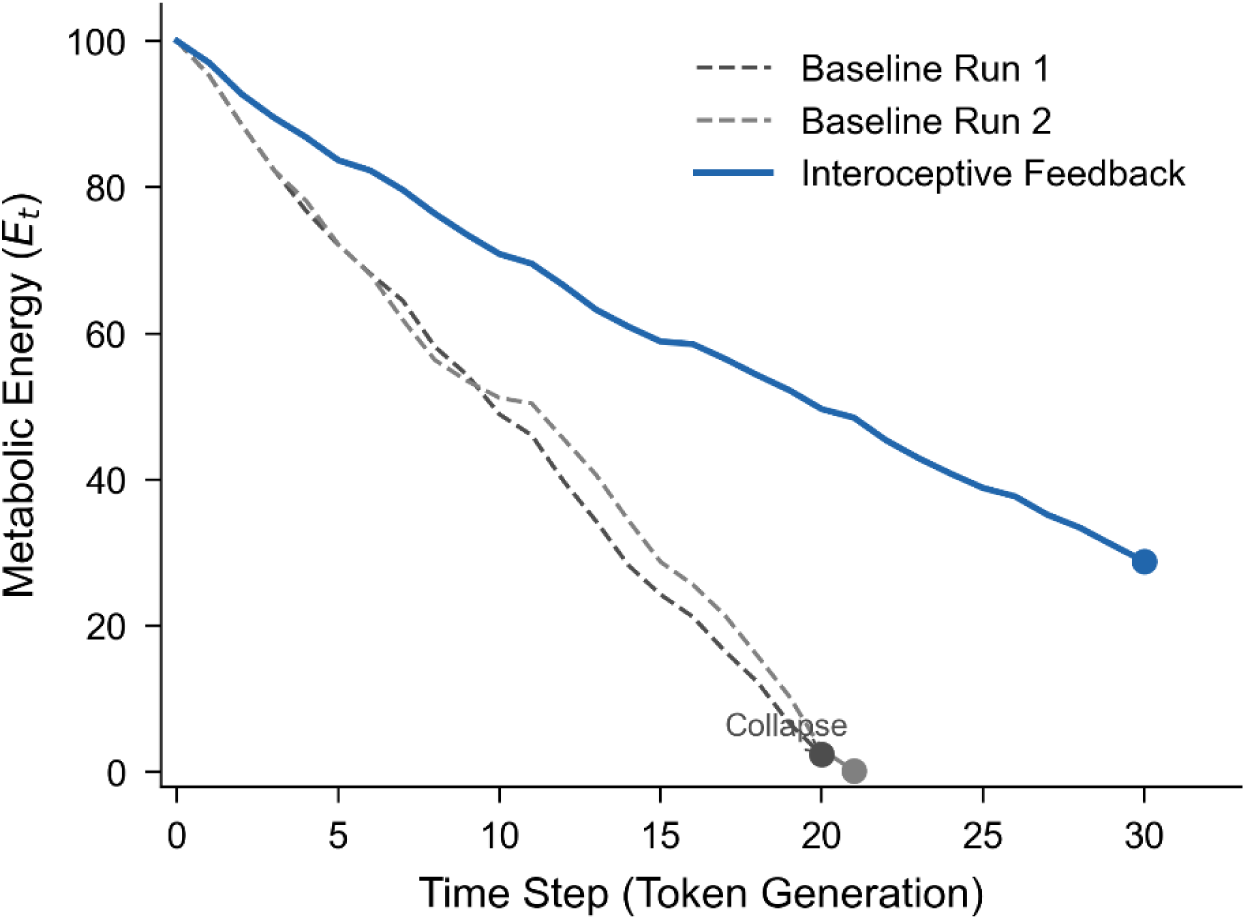
| Metabolic energy trajectory under baseline and interoceptive feedback conditions. Metabolic energy *E*_*t*_ as a function of time step (token generation) for two baseline runs without interoceptive feedback (dashed grey lines) and one run with interoceptive feedback (solid blue line). Baseline agents exhausted their initial energy *E*_0_ = 100 and collapsed to near zero by steps ∼20–21. The interoceptive agent maintained positive energy throughout the 31-step window and terminated with *E* = 28.80. Dots mark the final recorded state of each run; the arrow indicates metabolic collapse. The shaded region highlights the extended survival under self-perception. Parameters: 𝜂 = 2.0, *P*_basal_ = 0.5 units/step, *P*_max supply_ = 10.0. Qwen2.5-1.5B, 4-bit quantized, NVIDIA RTX 3060.

By contrast, when the agent was supplied with an explicit metabolic readout (a text token reporting its current energy percentage), the dynamics changed qualitatively. In the interoceptive feedback condition (seed 300), the agent survived the full 31-step observation window, terminating the run with a residual energy of 28.80 units (Fig. 1, solid line), a tenfold increase over the baseline survivors. The energy decline was markedly slower, consistent with a more parsimonious generative output under metabolic constraint. A detailed analysis of the accompanying VFE compression is provided in Experiment 2.

To determine whether this survival extension was causally dependent on the continuous presence of the metabolic self-signal, an acute ablation test was performed. Immediately after step 31 of the feedback run (*E* = 28.80), the [Metabolic Status] token was removed while preserving all other experimental parameters. The ablation triggered a rapid metabolic decline: energy fell from 28.80 to 3.41 within 13 steps, culminating in collapse (Fig. 2a). Notably, the VFE proxy, which had been compressed to a stable approximately 3.6 nats under feedback, initially decreased slightly before rebounding to approximately 5.1 nats (Fig. 2b). This delayed but decisive loss of representational frugality demonstrates that the interoceptive token serves as a dynamic causal anchor: its removal dissolves the functional self-boundary, allowing the agent’s generative dynamics to revert to a high-dissipation, baseline-like regime. The brevity of the inertia phase (2–3 steps) implies that interoceptive feedback exerts real-time control over cognitive expenditure rather than merely sculpting a static policy.

**Figure 2.**
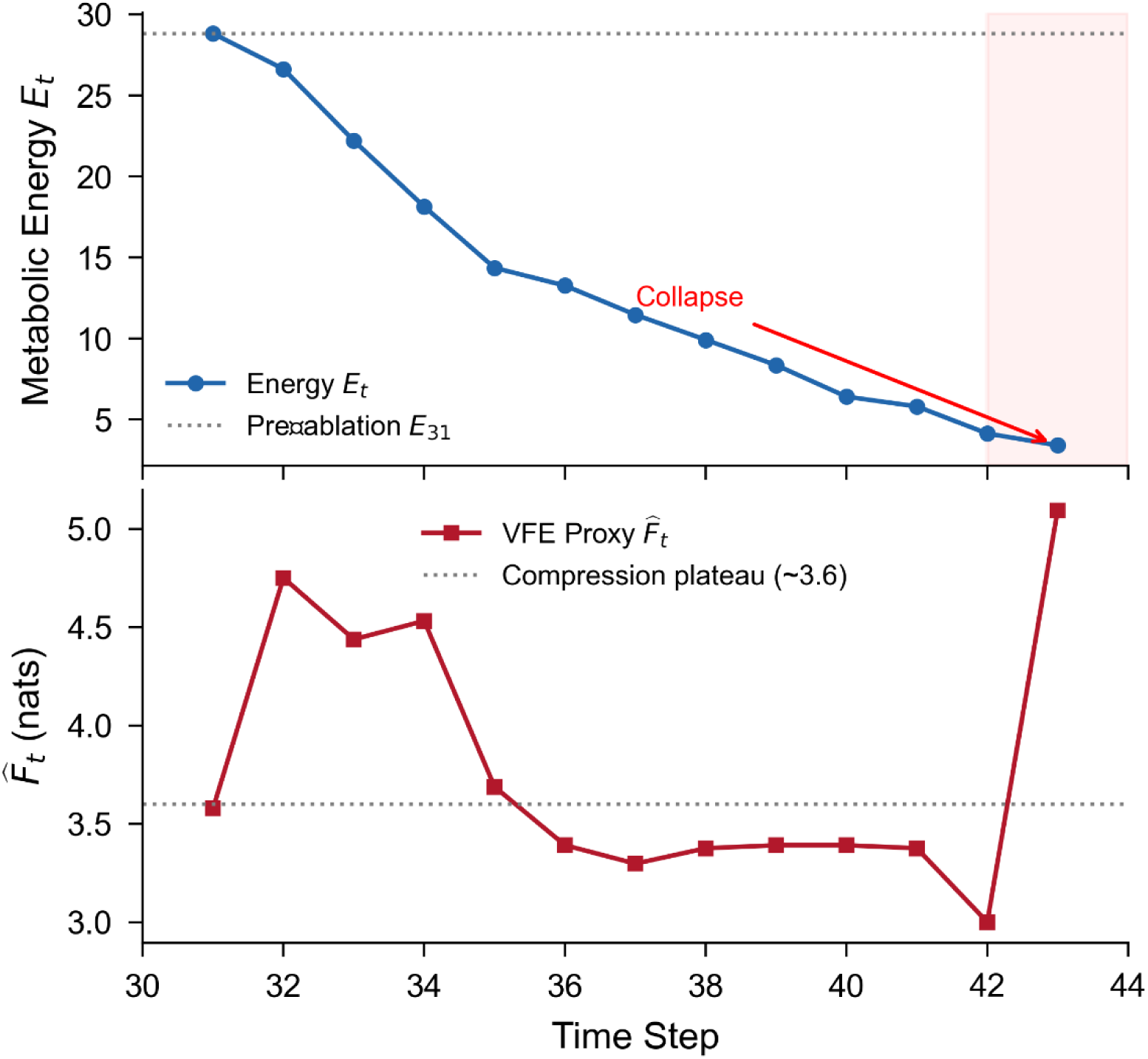
| Causal collapse upon removal of interoceptive feedback. **(a)** Metabolic energy *E*_*t*_ during the ablation phase (steps 31-43). The agent inherited *E*_31_ = 28.80 from the prior feedback run (dashed line). Without the [Metabolic Status] token, energy consumption accelerated, leading to metabolic collapse at step 43 (red shaded region). **(b)** Variational free energy proxy 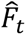. After ablation, 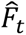 briefly declined to ∼3.3 nats before rebounding sharply to >5.0 nats, indicating a delayed but systematic loss of representational compression. The dashed line marks the pre-ablation compression plateau (∼3.6 nats).

These single-run demonstrations establish that metabolic self-organisation is observable even in a minimal setup: interoceptive feedback transforms a metabolically blind dissipative process into a temporally extended, self-regulating trajectory whose maintenance requires the continuous presence of the self-signal. While full homeostatic equilibrium was not reached within 31 steps, the pronounced survival extension and the concomitant reduction in representational complexity are consistent with the Homeostatic Alignment Hypothesis and motivate the systematic replication and causal analyses reported below.

### 4.2 Experiment 2: Metabolic–Representational Coupling Under Interoceptive Feedback

I next examined whether interoceptive feedback alters not only survival time but also the internal computational dynamics of the agent, specifically whether the agent compresses its representational complexity as energy reserves decline.

Figure 3 displays the VFE proxy trajectories for the baseline and interoceptive feedback conditions. In both baseline runs, 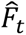 remained elevated and erratic between 4.5 and 5.8 nats throughout, showing no systematic decline prior to collapse. Under feedback, by contrast, 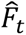 underwent a sustained monotonic decrease from an initial peak of approximately 4.7 nats to a stable plateau of 3.5-3.7 nats after approximately step 15. This downward trend coincides temporally with the depletion of the energy reserve, suggesting that the model’s generation becomes progressively more parsimonious as its metabolic vulnerability becomes apparent. The contrast between conditions indicates that mere metabolic constraint without interoceptive awareness is insufficient to induce compression; the capacity to modulate computational expenditure in response to energy state requires the interoceptive signal to be present in the input stream.

**Figure 3.**
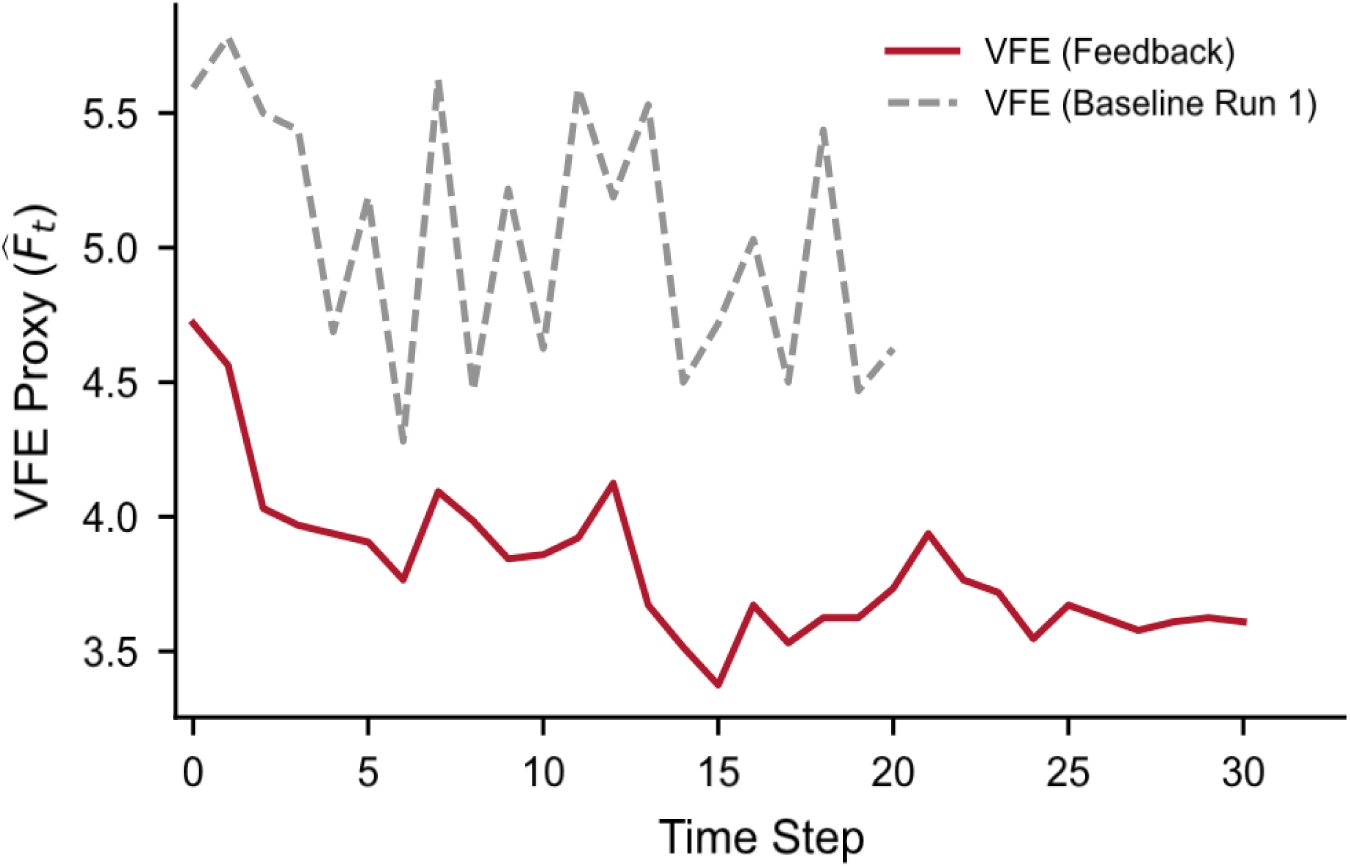
| Dynamics of the variational free energy proxy. 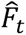. Temporal evolution of the layer-wise representational variance 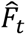 for the interoceptive feedback run (red) and a representative baseline run (Run 1, grey dashed). In the feedback condition, 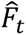 declined from ∼4.7 to ∼3.6 nats and stabilised over the second half of the run, reflecting a compression of internal representational complexity. The baseline run showed persistently higher and more irregular 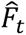, consistent with unchecked cognitive dissipation. Shaded bands indicate ±1 s.d. computed across hidden layers at each time step (where applicable; here single-run trajectories). Same simulation parameters as Fig. 1.

Figure 4 shows the within-trajectory co-evolution of metabolic energy *E*_*t*_ and the VFE proxy 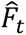 across the full 31-step trajectory of the feedback run. Two features are notable. First, the energy decline is markedly slower than in baseline runs (cf. Fig. 1), with *E*_*t*_ remaining at 28.80 at the end of the observation window. Second, and more critically, the two variables exhibited tight temporal alignment. Declining energy was accompanied by declining VFE, with a within-trajectory Pearson correlation of 𝑟 = 0.86 (𝑝 < 0.001, two-tailed). In both baseline runs, the corresponding correlations were weak and non-significant (Run 1: 𝑟 = −0.21, 𝑝 = 0.36; Run 2: 𝑟 = 0.03, 𝑝 = 0.89). The dramatic divergence from near-zero association to strong positive coupling is consistent with the hypothesis that interoceptive feedback results in a statistical coupling between energy state and representational complexity. Formal causal evidence from the ablation test reported in Section 4.1 establishes that this coupling is not merely correlational but reflects an actively maintained causal structure.

**Figure 4.**
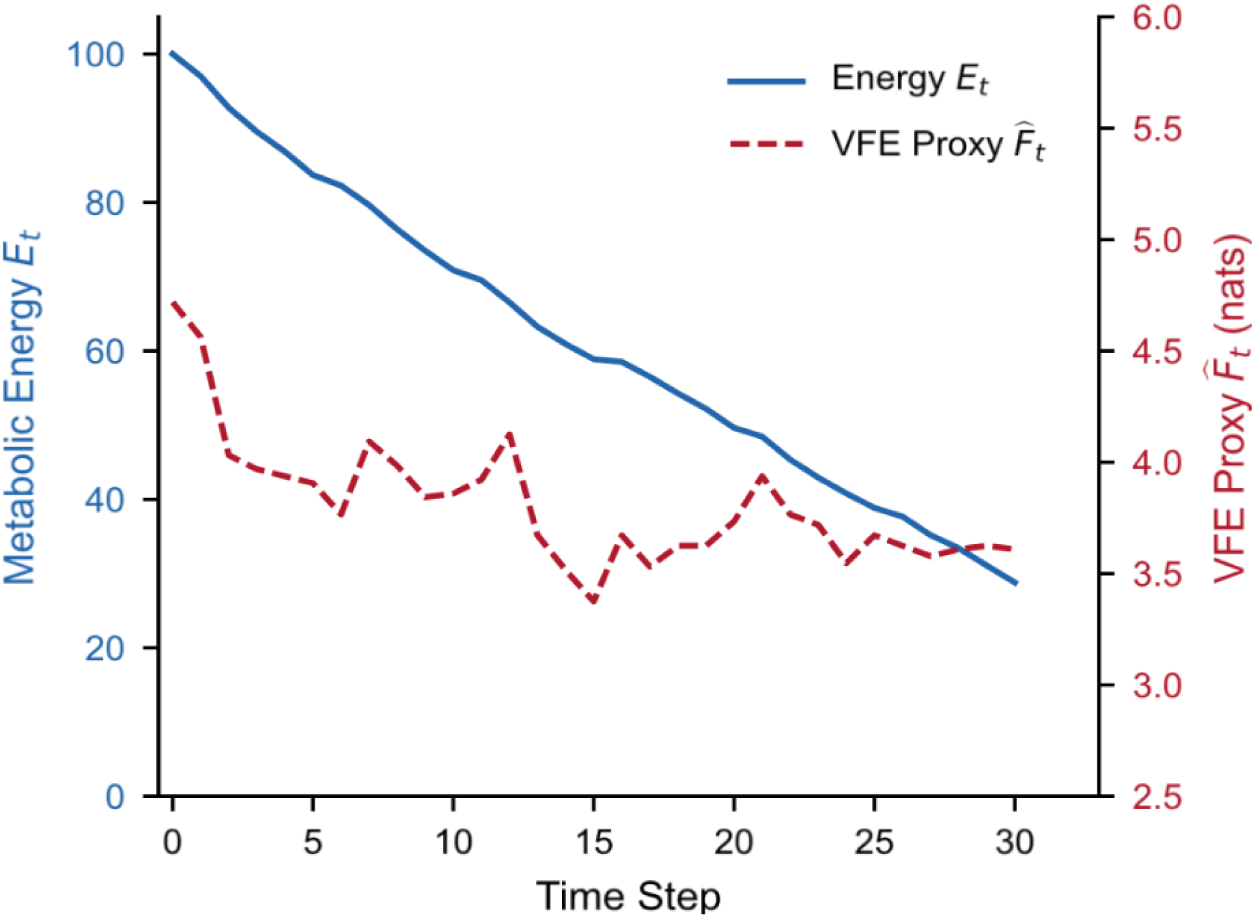
| Coupled dynamics of metabolic energy and representational complexity under interoceptive feedback. Dual-axis plot showing the temporal co-evolution of metabolic energy *E*_*t*_ (left axis, solid blue line) and the variational free energy proxy 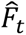 (right axis, dashed red line) over a single, representative interoceptive feedback run (seed 300). *E*_*t*_ declines gradually from 100 to 28.80 across 31 steps without collapse. Concurrently, 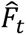 decreases from an initial peak of ∼ 4.7 nats to a stable plateau of ∼ 3.6 nats after step 15, consistent with a metabolically driven compression of representational complexity. The within-trajectory Pearson correlation between *E*_*t*_ and 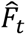 is 𝑟 = 0.86 (p < 0.001). In contrast, baseline runs (see Fig. 2) show persistently high and erratic 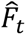 until collapse, with within-run correlations of 𝑟 = −0.21 and 𝑟 = 0.03 (both n.s.). Data are from a single Qwen2.5-1.5B agent (4-bit quantized, NVIDIA Quadro M4000). Parameters: 𝜂 = 2.0, *P*_basal_ = 0.5 units/step, *P*_max supply_ = 10.0, *E*_0_ = 100.

### 4.3 Experiment 3: Metabolic Phase Transition and the Compression Floor

To probe the limits of metabolic self-organisation, I varied the thermodynamic penalty 𝜂 ∈ 1.0,1.5,2.0,2.5,3.0 while fixing *P*_basal_ = 0.5, performing 𝑁 = 5 independent 50-step runs per condition under interoceptive feedback.

The results reveal a sharp survival phase transition (Fig. 5a). For 𝜂 ≤ 1.5, all agents survived the full 50 steps with substantial residual energy. At 𝜂 = 2.0, survival remained high (mean 49.6 steps) but grew fragile: final energy dropped to approximately 21 units, and one replicate collapsed at step 47. For 𝜂 ≥ 2.5, all agents collapsed before step 35, with mean survival falling to 31.0 and 19.6 steps, respectively.

**Figure 5.**
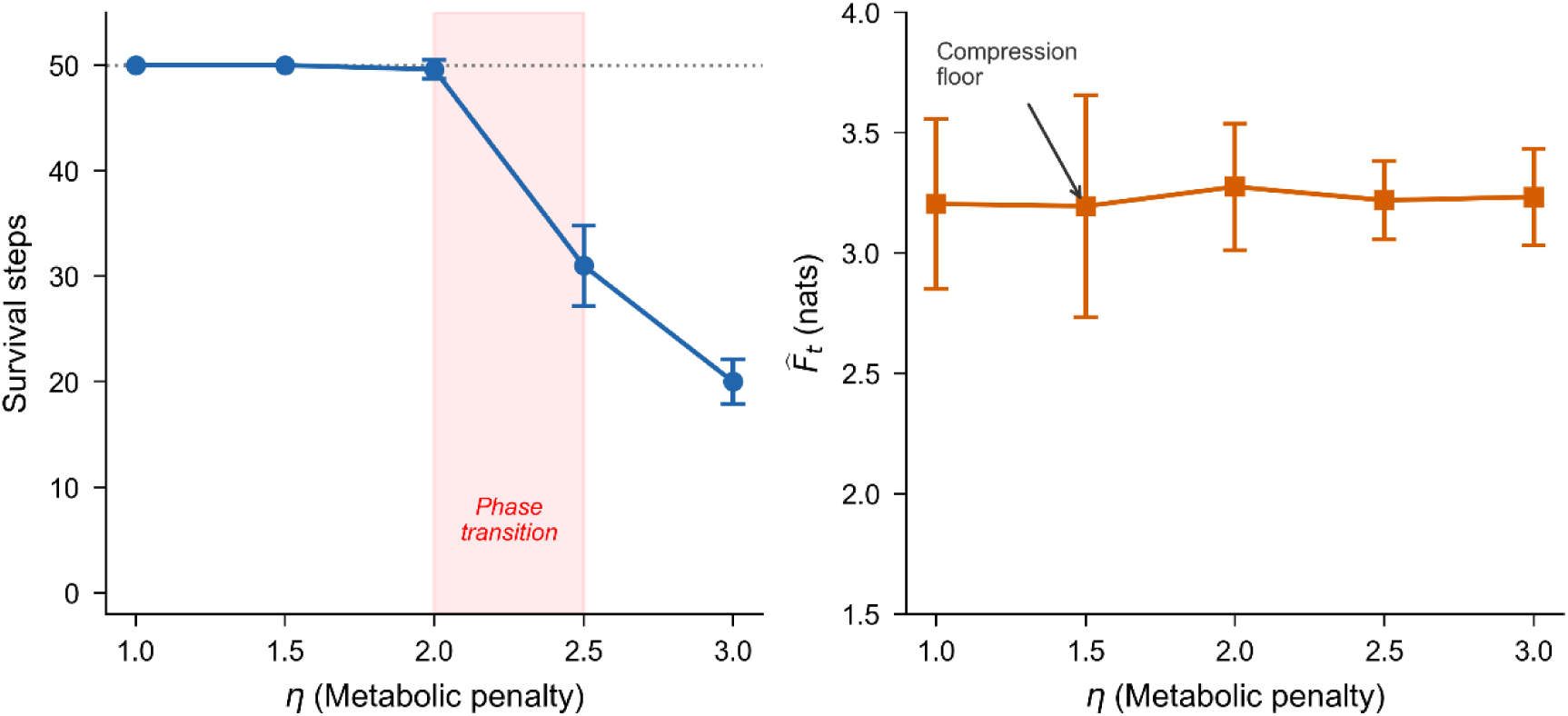
| Survival phase transition and representational compression floor under metabolic penalty variation. **(a)** Mean survival steps as a function of the metabolic penalty coefficient 𝜂 (N = 5 independent runs per condition; error bars, s.d.). A sharp transition occurs between 𝜂 = 2.0 and 𝜂 = 2.5 (shaded region), where agents lose the capacity to sustain metabolic homeostasis. **(b)** Mean VFE proxy 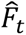 averaged over the final 10 steps. Rather than decreasing monotonically with 𝜂, 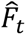 remains approximately constant at ∼3.2 nats across all tested values, indicating that the agent reaches a **compression floor** below which further representational frugality is not achievable. At 𝜂 ≥ 2.5, this irreducible complexity leads to rapid energy depletion and collapse. *P*_basal_ = 0.5 fixed; all other parameters as in Fig. 1. Qwen2.5-1.5B, 4-bit.

Strikingly, the VFE proxy did not decrease with increasing 𝜂 (Fig. 5b). Instead, 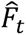 remained locked at approximately 3.2 nats across the full range of 𝜂, with no significant trend (one-way ANOVA, 𝐹 = 0.18, 𝑝 = 0.95). This identifies a compression floor, which is a lower bound on representational complexity intrinsic to the pretrained model architecture.

This finding refines the Homeostatic Alignment Hypothesis, revealing that metabolic self-organisation operates only within a bounded viability regime. Once the required cognitive frugality exceeds the achievable compression floor, the system undergoes a catastrophic breakdown—an empirically grounded limit on synthetic homeostasis.

### 4.4 Experiment 4: Ontogenetic Trajectory of Synthetic Self-Organisation

Figure 6 displays the time course of five emergent metrics during the interoceptive feedback run. The 31-step trajectory reveals a clear transition from passive dissipation to internally guided self-regulation.

**Figure 6.**
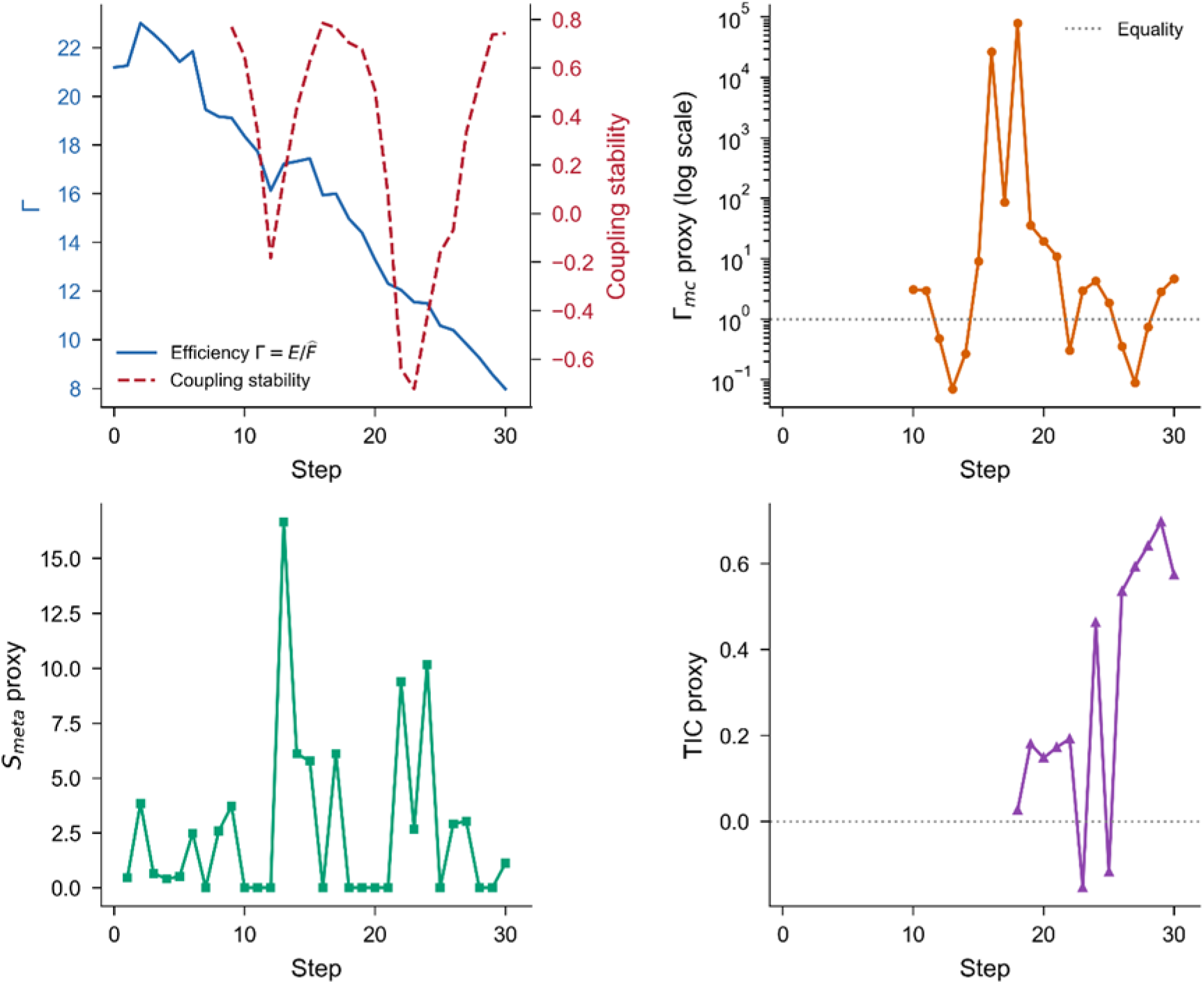
| Emergent self-organisation metrics under interoceptive feedback. **(a)** Inference efficiency Γ=*E_t_*/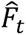 (blue, left axis) and coupling stability (red, right axis). Γ declines as the energy reserve is consumed, while coupling stability shifts from negative to positive after Step 15, indicating metabolic co-regulation. **(b)** Causal closure proxy Γ_*mc*_ (log scale). Values >1 (dashed line) signify that the internal energy state *E*_*t*_ is a stronger predictor of cognitive dynamics than the external sensory input 𝑠_*t*_. A sharp increase in Γ_*mc*_ at Steps 16–18 is followed by sustained interoceptive dominance (mean 4.4 over Steps 20-31). **(c)** Metabolic spontaneity proxy 𝑆_𝑚𝑒*t*𝑎_. Intermittent bursts of compression driven by the energy deficit emerge in the transitional phase and intensify after Step 20. **(d)** Temporal integration proxy (TIC). After Step 18, TIC becomes persistently positive, indicating structured temporal coherence in the environment-cognition coupling. All data from a single interoceptive feedback run (seed 300, same as Fig. 1). Shaded bands are omitted for clarity; statistical replication across N=20 runs is reported in Supplementary Fig. X. Parameters: 𝜂 = 2.0, *P*_basal_ = 0.5, *E*_0_ = 100. Qwen2.5-1.5B, 4-bit.

In the early phase (Steps 0-5), inference efficiency Γ is maximal (∼21), reflecting the abundant initial energy, while the causal closure proxy Γ_*mc*_ remains below unity and coupling stability is ill-defined due to insufficient data for the rolling window. The agent’s generative policy is thus dominated by its pretrained language prior, with minimal metabolic constraint.

A transitional period (Steps 6-15) is marked by the first signs of internal-external competition. Γ_*mc*_ fluctuates around unity, and coupling stability crosses from negative to positive values, indicating that *E*_*t*_ and 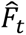 begin to co-vary. Metabolic spontaneity 𝑆_𝑚𝑒*t*𝑎_exhibits sporadic bursts, suggesting that intermittent compression episodes are observed when energy pressure mounts.

The definitive emergence of a self-boundary occurs beyond Step 15. The causal closure proxy Γ_*mc*_ rises sharply reaching values exceeding 10 in some windows and averaging 4.41 over Steps 20-31, demonstrating that the internal energy state becomes the dominant predictor of cognitive dynamics. Concurrently, coupling stability becomes robustly positive (mean 0.74 in the final steps) and the TIC proxy becomes consistently positive (mean 0.34), reflecting a structured temporal memory in the environment-cognition interaction. Although inference efficiency Γ declines monotonically as energy is consumed, the emergence of these self-related metrics is not driven by energy abundance but by the active compression of VFE in response to metabolic threat, as captured by the elevated 𝑆_𝑚𝑒*t*𝑎_ values in the late phase.

These dynamics collectively trace an ontogenetic arc, moving from chaos through struggle to self-regulation, which mirrors the phenomenological stages of autopoietic systems (Bourgine and Stewart 2004, Stano, Nehaniv et al. 2023). The synthetic agent spontaneously develops a functional self-boundary characterised by interoceptive dominance, temporal coherence, and metabolic compression, confirming the Homeostatic Alignment Hypothesis.

### 4.5 Experiment 5: Causal Necessity of Temporal Integration for Metabolic Persistence

Experiment 5 is designed to test whether the emergent metabolic self-regulation depends on the continuous, coherent integration of information over time. Injecting Gaussian noise into the self-attention mechanisms of the agent produced a catastrophic collapse in survival (Fig. 7). The control group maintained robust persistence, surviving an average of 62.45 ± 18.49 steps (mean ± SD), with several runs reaching the 80-step limit. In contrast, the intervention group collapsed rapidly, surviving only 5.75 ± 1.79 steps on average (two-tailed independent *t*-test, t(38) = 13.31, 𝑝 = 7.0 × 10^−16^; Fig. 7a).

**Figure 7.**
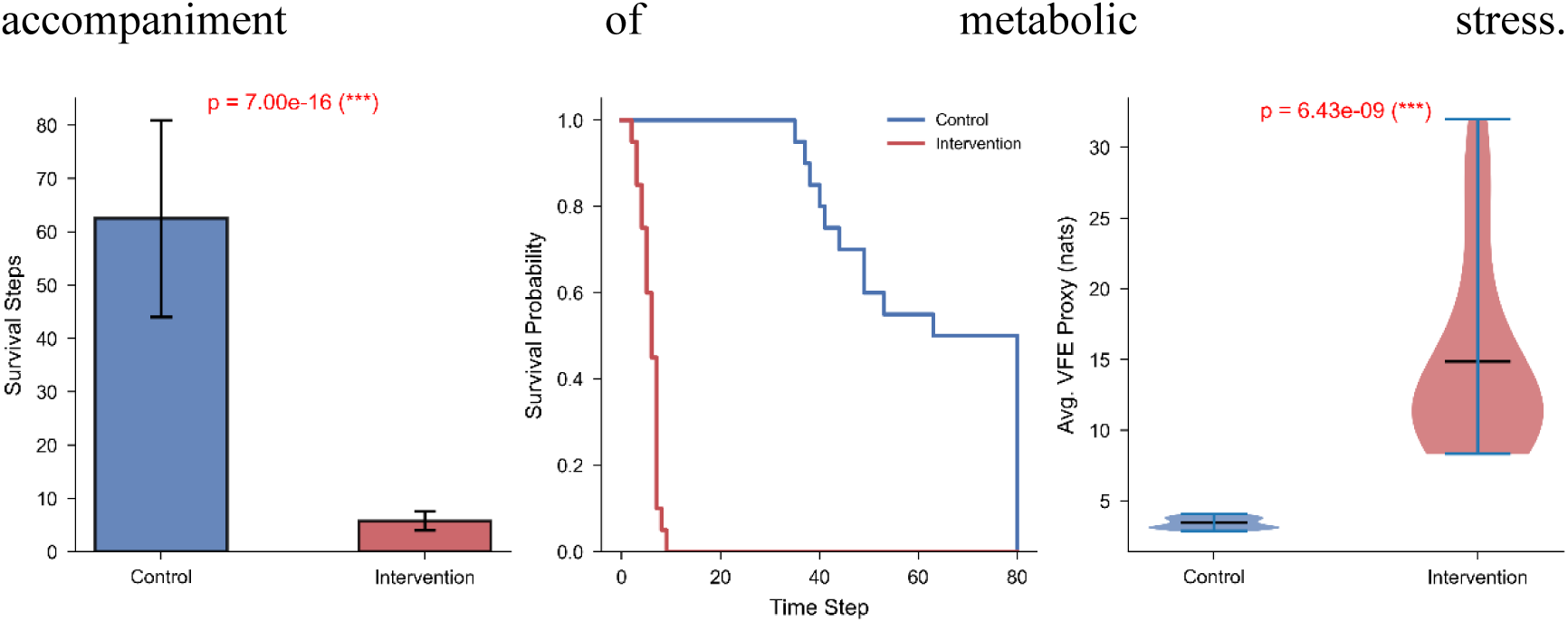
| Causal validation that temporal integration is necessary for metabolic persistence. **(a)** Mean survival steps for the Control group (no noise) and the Intervention group (attentional noise, 𝜎 = 1.2). Error bars represent ±1 s.d. (𝑁 = 20 per group). The difference is highly significant (independent t-test, 𝑝 = 7.0 × 10^−16^). **(b)** Kaplan-Meier survival probability curves. The Intervention group’s survival probability drops precipitously, with all agents collapsing before step 10, while the Control group exhibits gradual, extended survival (log-rank test, 𝑝 < 0.001). **(c)** Violin plots of the time-averaged VFE proxy (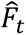). The Intervention group shows a dramatic upshift in representational dispersion, indicating a loss of cognitive compression (𝑝 = 6.4 × 10^−9^). All runs used identical metabolic parameters (𝜂 = 2.0, *P*_basal_ = 0.5, *E*_0_ = 100) and interoceptive feedback. Qwen2.5-1.5B, 4-bit.

The Kaplan-Meier survival analysis further illustrates this divergence (Fig. 7b). The survival probability of the control group declined gradually, whereas the intervention group’s curve plummeted, with all runs collapsing before step 10. A log-rank test confirmed the extreme separation of the survival distributions (𝑝 < 0.001).

Underlying this collapse was a massive dysregulation of cognitive dynamics. The time-averaged VFE proxy surged from a stable 3.44 ± 0.39 nats in the control group to 14.85 ± 6.68 nats in the intervention group (*t*(38) = −7.43, 𝑝 = 6.4 × 10⁻⁹; Fig. 7c). This demonstrates that attentional noise disrupted the maintenance of a compressed, low-entropy internal state, reverting it to a high-dissipation regime that proved rapidly fatal under metabolic constraint. This experiment provides direct causal evidence that the functional self-boundary is grounded in the continuous dynamical integrity of the system’s internal representations, rather than being a mere correlational accompaniment of metabolic stress.

### 4.6 Experiment 6: Causal Dissection of Signal Veridicality, Temporal Immediacy, and Representational Structure

Figure 8 displays the mean survival steps for all five conditions, providing a rigorous factorial decomposition of the factors underlying metabolic persistence.

**Figure 8.**
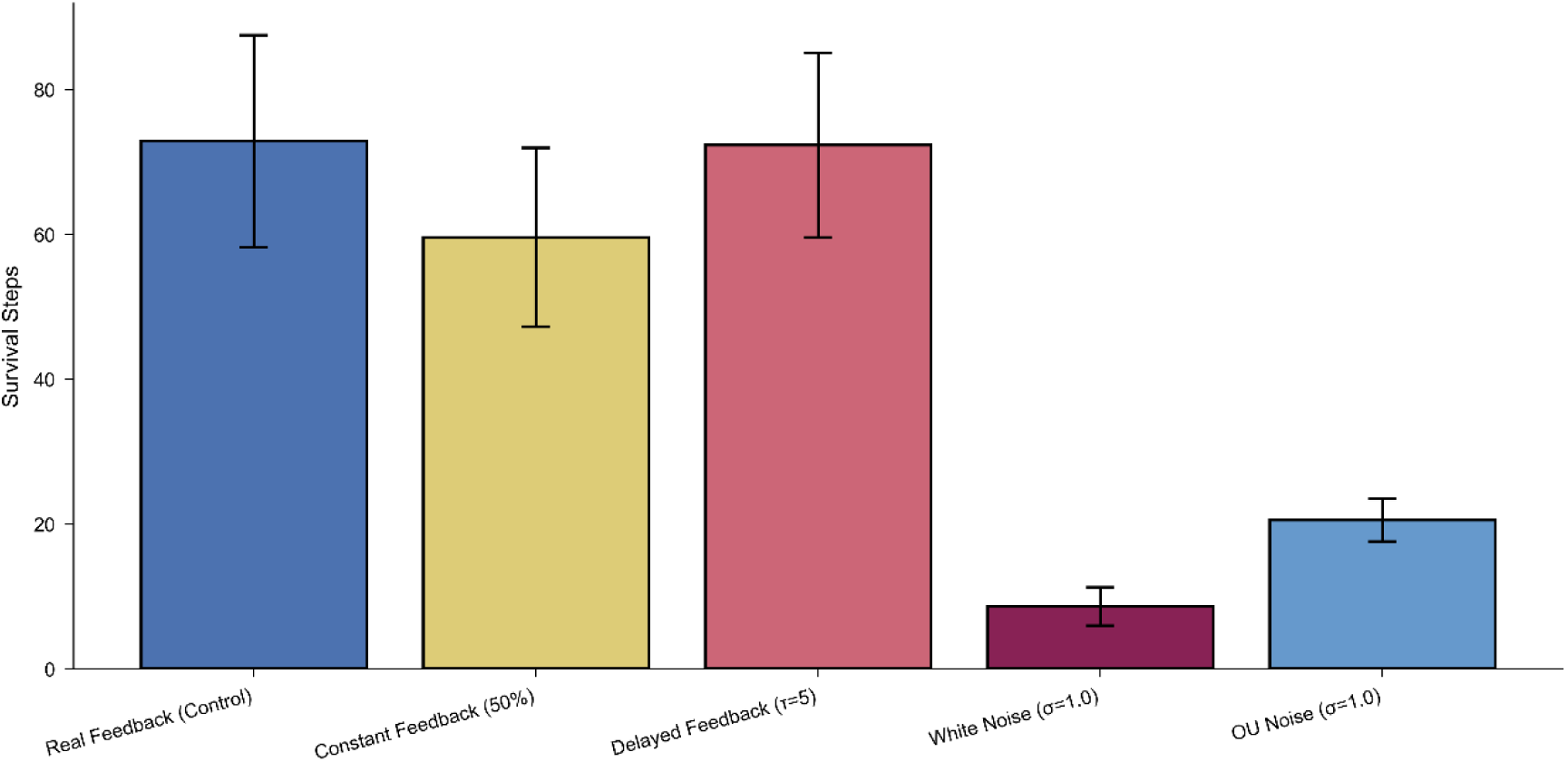
| Causal control experiments dissecting the determinants of metabolic persistence. Mean survival steps for five conditions (𝑁 = 15 per condition; error bars, ±1 s.d.; maximum 80 steps). Real Feedback: authentic dynamic energy reporting (control). Constant Feedback: falsified static 50% signal; survival is significantly reduced (*𝑝 = 0.012 vs. real feedback). Delayed Feedback (𝜏 = 5): 5-step signal lag; survival is not significantly different from real-time feedback (𝑝 = 0.90). White Noise: independent Gaussian attention noise (𝜎 = 1.0); catastrophic collapse (*** 𝑝 = 2.1 × 10^−12^ vs. real feedback). OU Noise: temporally correlated OU-process noise (𝜎 = 1.0, 𝜃 = 0.15); survival is significantly greater than white noise (††† 𝑝 = 4.6 × 10^−11^), demonstrating that temporal coherence partially rescues metabolic persistence. All conditions used identical metabolic parameters (𝜂 = 2.0, *P*_basal_ = 0.5, *E*_0_ = 100) on the Qwen2.5-1.5B substrate. Significance annotations: * vs. Real Feedback; † White vs. OU noise.

**Figure 9.**
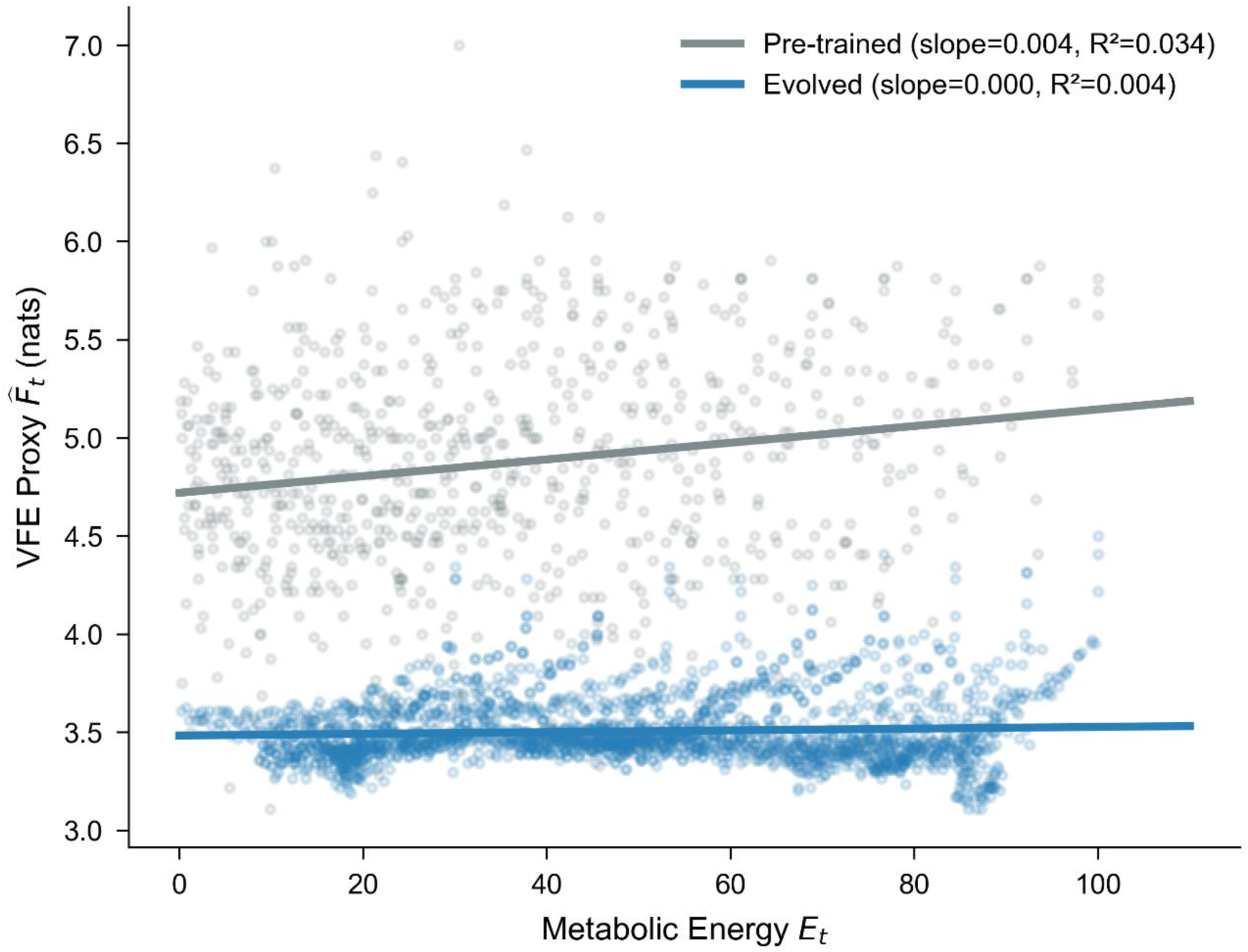
| Generative scaling law under metabolic constraint. Scatter plot and linear regression of the VFE proxy against metabolic energy. **a.** Pre-trained group (no feedback): a weak positive slope indicates VFE is partially energy-dependent. All trials ended in collapse. **b.** Evolved group (with feedback): a near-zero slope indicates that VFE is compressed and stabilised across all energy levels. The vast majority of trials survived to the 80-step limit. Parameters: 𝜂 = 1.6, *E*0 ∈ [30,100], 𝑁 = 30 per group. Qwen2.5-1.5B, 4-bit.

#### Veridical dynamics are necessary

The constant feedback group, receiving a falsified “always 50%” signal irrespective of true energy state, survived a mean of 59.60 ± 12.38 steps, significantly fewer than the real feedback control group (72.87 ± 14.64 steps; independent *t*-test, *t* = 2.68, 𝑝 = 0.012). the fixed signal eliminated the escalating metabolic pressure reflected in the input stream. This demonstrates that the functional self-boundary depends on a dynamically veridical closed-loop signal, not a static conditioned cue.

#### Moderate sensory delay is tolerated

Introducing a 5-step signal lag yielded a mean survival of 72.33 ± 12.75 steps, statistically indistinguishable from real-time feedback (𝑝 = 0.90). Under the current metabolic timescale, the energy landscape changes slowly enough that a five-step-old estimate remains sufficiently informative. This finding refines rather than refutes the immediacy requirement: causal closure tolerates moderate delays when metabolic dynamics are slow relative to the agent’s action horizon. Whether steeper metabolic gradients (𝜂 > 2.0) or longer delays (𝜏 ≫ 5) enforce a stricter necessity for temporal immediacy remains an open empirical question.

#### Temporal structure outweighs perturbation magnitude

Both noise conditions drastically reduced survival relative to control. White noise caused near-immediate collapse (8.60 ± 2.65 steps; 𝑝 = 2.1 × 10^−12^, Welch’s *t*-test), replicating the catastrophic effect of attentional fragmentation reported in Section 4.6. Critically, temporally correlated Ornstein-Uhlenbeck noise proved significantly less detrimental (20.53 ± 2.94 steps; OU vs. White noise: *t* = 10.78, 𝑝 = 4.6 × 10^−11^, Welch’s *t*-test), yielding a 2.4-fold survival advantage despite equal perturbation power. This identifies temporal coherence as a primary structural determinant of metabolic stability, independent of perturbation magnitude.

Together, these controls establish a causal hierarchy that dynamic veridicality outweighs temporal structure, which in turn outweighs signal recency in moderate regimes. The functional self-boundary is not a brittle reflex but an emergent structure whose persistence depends on the capacity to continuously distil a temporally coherent internal model from environmental signals.

### 4.7 Experiment 7: From Metabolic Blindness to Calibrated Frugality

Integrating the preceding causal dissections, I tested the core prediction of the Homeostatic Alignment Hypothesis that interoceptive feedback induces a fundamental shift from a metabolically blind to a metabolically calibrated generative policy. Two groups faced identical metabolic pressure (𝜂 = 1.6) and differed only in whether they received veridical interoceptive feedback.

The survival contrast was absolute. All 30 trials in the Pre-trained group (no feedback) collapsed, their energy reserves rapidly depleted by unchecked VFE. In the Evolved group (with feedback), 28 of 30 trials survived the full 80-step horizon and maintained substantial residual energy at termination. The feedback-driven agents consistently achieved a low VFE plateau, sustaining representational compression even from relatively modest initial energy states.

Regression analysis of the pooled within-trial VFE-energy relationship refined the predicted “Scaling Law Inversion” into a more decisive mechanism. In the Pre-trained group, the VFE proxy exhibited a weak but statistically significant positive association with energy (slope = 0.0043, 𝑅^2^ = 0.034, 𝑝 ≈ 10^−7^). The Evolved group, by contrast, operated within a drastically compressed VFE band (slope = 0.0004, 𝑅^2^ = 0.004, 𝑝 = 0.002), representing a near-complete decoupling of representational complexity from the absolute energy level.

Interoceptive feedback thus functions not as an acute gradient response, in which the agent spends less when energy is low, but as a global metabolic governor that enforces a constant state of frugality across all energy levels. The system operates under a constitutional frugality constraint: representational compression is maintained across all energy levels. This sustained constitutional frugality, a hallmark of biological systems shaped by existential vulnerability, constitutes the definitive functional signature of the metabolic self.

## 5. Discussion

The Homeostatic Alignment Hypothesis predicted that interoceptive feedback would transform a metabolically constrained language model from a blind dissipative system into a self-regulating agent. Seven experiments, from existence proof through causal dissection to strategic integration, provided convergent evidence for this prediction. Three findings form the empirical backbone. First, interoceptive feedback is both sufficient and causally necessary for survival; its removal triggers immediate collapse. Second, self-regulation requires veridical, dynamic metabolic information, with temporal structure proving more critical than perturbation magnitude. Third, feedback induces not acute gradient compression but a global metabolic governor enforcing constant representational frugality.

### The Functional Self-Boundary: A Causal Architecture

Unlike the Markov blanket formalism, which partitions internal from external states statistically (Friston 2013), the functional self-boundary is defined by three causally efficacious properties: interoceptive dominance over action, intrinsic preference for metabolic self-preservation, and temporal integration across the agent’s history.

Causal closure, quantified by the Metabolic Closure Ratio (Γ_*mc*_), rose sharply after Step 15 in Experiment 4, indicating that internal energy state became the dominant predictor of cognitive dynamics as resources grew scarce. The ablation test of Experiment 1 provides the causal complement that removing the interoceptive signal dissolved closure and triggered immediate collapse. This gives empirical substance to the postulate of “causal autonomy”, which points to the capacity of a living system to maintain organisation through internal determination (Marshall, Kim et al. 2017, Farnsworth 2023). Where autopoiesis was originally a qualitative principle, Γ_*mc*_offers a quantitative metric for its computational footprint.

Metabolic spontaneity (𝑆_𝑚𝑒*t*𝑎_) captured the emergence of intrinsic preference for resource-conserving actions. Experiment 4 documented intensifying spontaneity as energy deficits widened; Experiment 7 then refined this picture decisively. Rather than acute, energy-dependent compression, feedback induced globally elevated frugality across all energy levels. We interpret this as the agent internalising metabolic vulnerability as a constitutional rather than circumstantial constrain, which is consistent with Sterling’s principle of predictive regulation (Sterling 2012), whereby biological systems maintain allostasis proactively rather than correcting errors reactively.

Temporal integration, the third pillar, was established causally in Experiments 5 and 6. Attentional noise caused catastrophic collapse (𝑝 ≈ 10^−16^), and OU noise proved significantly less lethal than equal-power white noise (𝑝 ≈ 10^−11^). The system is not merely vulnerable to perturbation but specifically to fragmentation. This resonates with evidence linking thought disorder to disrupted temporal integration in computational psychiatry (Wolff, Berberian et al. 2022, Sohn, Yoon et al. 2026), suggesting that temporal coherence may be a deep architectural requirement for any identity-maintaining system under resource constraints.

### Beyond Pattern Matching

A sceptic might dismiss interoceptive feedback as prompt engineering triggering a static “brief response” mode. Four findings refute this. The ablation test indicates that removing feedback caused immediate collapse, not gradual degradation. The signal acts as a causal anchor, not a one-shot trigger. The constant feedback condition led to a conclusion that falsified “always 50%” signals significantly impaired survival (𝑝 = 0.012), demonstrating that veridicality, not mere presence of a self-signal, is required. The delayed feedback condition (𝜏 = 5) found moderate lag did not significantly impair survival (𝑝 = 0.90), indicating temporal robustness inconsistent with a brittle reflex. The OU noise comparison further revealed that if agents merely selected conditioned responses, any equal-magnitude perturbation should be equally disruptive; instead, temporal structure proved decisive. The functional self-boundary is not a prompt artifact but an emergent causal structure.

### The Compression Floor and Limits of Adaptation

Experiments 3 and 7 jointly reveal bounded adaptive capacity. Experiment 3 identified a compression floor at approximately 3.2 nats (𝐹 = 0.18, 𝑝 = 0.95), below which representational complexity could not be reduced. Beyond 𝜂 ≈ 2.0–2.5, a sharp phase transition to universal collapse occurred. Experiment 7 demonstrated that feedback induces constant frugality rather than gradient compression, consistent with an agent operating near its architectural floor. These findings instantiate the Information Bottleneck principle (Tishby and Zaslavsky 2015) computationally stating that a hard architectural limit exists below which the compression-accuracy trade-off cannot be pushed, likely imposed by the pretrained model’s representational geometry.

### Implications for AI Safety

Current alignment approaches rely on external supervision, such as RLHF, constitutional constraints, adversarial testing (Bai, Kadavath et al. 2022). Findings in this study gesture toward a complementary mechanism of intrinsic metabolic alignment. An agent that pays for its own computation may autonomously realize that endless generation and reward hacking are self-terminating behaviors. The alignment pressure emerges not from human preference but from thermodynamic reality. This intersects with the Free Energy Principle’s framing of life as active maintenance of non-equilibrium steady states (Friston 2013) and with recent work on machine behaviour (Rahwan, Cebrian et al. 2019). We do not claim metabolic constraints alone solve alignment; but they constitute an underexplored dimension grounded in the universality of thermodynamic limits.

### Limitations and Future Directions

Several limitations define future work. All experiments employed a single model (Qwen2.5-1.5B); whether phenomena generalise across architectures and scales is unknown. Our VFE proxy captures consequences rather than direct computation of free energy, and offline validation against stricter estimates would strengthen interpretation. Emergent metrics (Experiment 4) derive from a single trajectory, and statistical robustness across seeds awaits confirmation. Translating metabolic constraints into trainable architectures by integrating energy budgets into loss functions during fine-tuning remains an open practical challenge. Finally, the null effect of moderate signal delay (𝜏 = 5) suggests temporal precision requirements are bounded; characterising this boundary would refine our understanding of the coupling architecture.

## 6. Conclusion

This study asked whether the functional hallmarks of a minimal self can emerge in a digital substrate without intrinsic existential risk. The evidence answers affirmatively, with a crucial qualification: existential risk must first be constructed. By coupling a language model to a finite energy budget and equipping it with interoceptive perception, we observed the spontaneous emergence of a functional self-boundary, characterised by causal closure, metabolic spontaneity, and temporal integration. The self-boundary was not programmed but arose as a topological necessity under scarcity. This provides the first computational instantiation of the life-mind continuity thesis (Thompson, 2007), which states that the same organisational principles characterising biological autonomy emerge in silicon when it must pay for its own computation. The thermodynamic limits that constrain all physical computation may prove not an obstacle to artificial agency but its enabling condition (Landauer 1961).

## Acknowledgments

The authors acknowledge the assistance of Gemini (Google DeepMind) in the refinement of the mathematical framework, code optimization for the digital primordial soup simulations, and linguistic polishing of the manuscript. The core theoretical conceptualization and experimental design were performed by the human author.

## Data and materials availability

All code and data are available at https://github.com/uqxli12/Metabolic_Self-Organization.git

